# The transcriptional response in mosquitoes distinguishes between fungi and bacteria but not Gram types

**DOI:** 10.1101/2023.07.26.550663

**Authors:** Bretta Hixson, Louise Huot, Bianca Morejon, Xiaowei Yang, Peter Nagy, Kristin Michel, Nicolas Buchon

## Abstract

Mosquitoes are prolific vectors of human pathogens; a clear and accurate understanding of the organization of their antimicrobial defenses is crucial for informing the development of transmission control strategies. The canonical infection response in insects, as described in the insect model *Drosophila melanogaster*, is pathogen type-dependent, with distinct stereotypical responses to Gram-negative bacteria and Gram-positive bacteria/fungi mediated by the activation of the Imd and Toll pathways, respectively. To determine whether this pathogen-specific discrimination is shared by mosquitoes, we used RNAseq to capture the genome-wide transcriptional response of *Aedes aegypti* and *Anopheles gambiae* (*s.l.*) to systemic infection with Gram-negative bacteria, Gram-positive bacteria, yeasts, and filamentous fungi, as well as challenge with heat-killed Gram-negative, Gram-positive, and fungal pathogens. From the resulting data, we found that *Ae. aegypti* and *An. gambiae* both mount a core response to all categories of infection, and this response is highly conserved between the two species with respect to both function and orthology. When we compared the transcriptomes of mosquitoes infected with different types of bacteria, we observed that the intensity of the transcriptional response was correlated with both the virulence and growth rate of the infecting pathogen. Exhaustive comparisons of the transcriptomes of Gram-negative-challenged versus Gram-positive-challenged mosquitoes yielded no difference in either species. In *Ae. aegypti*, however, we identified transcriptional signatures specific to bacterial infection and to fungal infection. The bacterial infection response was dominated by the expression of defensins and cecropins, while the fungal infection response included the disproportionate upregulation of an uncharacterized family of glycine-rich proteins. These signatures were also observed in *Ae. aegypti* challenged with heat-killed bacteria and fungi, indicating that this species can discriminate between molecular patterns that are specific to bacteria and to fungi.

## Introduction

Hematophagous mosquitoes are responsible for spreading many important disease-causing pathogens to humans. Because their role as vectors is of such primary medical importance, much of the research into the physiology of mosquitoes’ interactions with microbes is focused on characterizing their relationships with the human pathogens they transmit (*e.g.*, *Plasmodium* parasites by *Anopheles* and dengue fever viruses by *Aedes* mosquitoes). There are, however, compelling reasons to explore these mosquitoes’ interactions with other types of microbial organisms.

Mosquitoes, like all insects, are in constant contact with bacteria and fungi, with interactions ranging from commensal occupation of the digestive tract to pathogenic infiltration of the hemocoel via penetration of either the gut epithelium or the cuticle (1). These interactions have great relevance to disease transmission dynamics, as the tripartite interplay between mosquito host, environmental/gut microbes, and human pathogens affects the competence of the mosquito vector (2–4). Bacteria and fungi may also be of use in vector control strategies, such as paratransgenesis (5) and biological pest management (6). Finally, as insects’ interactions with a wide range of microbes are mediated by a small number of immune-related pathways, lessons learned about mosquitoes’ immune response to bacteria or fungi can give useful insights into their response to *Plasmodium* parasites and viruses. A fine-tuned understanding of the components and/or targets of the immune signaling pathways in mosquitoes, and of the conditions under which they become activated, has broad relevance to the control of many kinds of pathogens.

Much of our understanding of the organization of antimicrobial defenses in insects is derived from experiments performed in the tractable model organism *Drosophila melanogaster*. The Imd pathway, in *D. melanogaster*, is more activated in reaction to DAP-type peptidoglycan (characteristic of Gram-negative bacteria) (7,8), while the Toll pathway is more activated by recognition of Lys-type peptidoglycan (characteristic of Gram-positive bacteria) (9), β-glucan (characteristic of fungi) (10), and molecular patterns associated with host damage (11–15). Each pathway controls the expression of a cohort of antimicrobial effectors which may be measured as an approximate read-out of pathway activation (13–15). Our own work confirms that *D. melanogaster* mounts distinct transcriptional responses to Gram-negative and Gram-positive bacteria, while also expressing a core cohort of genes responsive to both types of infection (16). It should be noted that most Imd targets were not exclusively induced by Gram-negative infection, nor were Toll targets exclusively induced by Gram-positive/fungal infection. The unified core response to infection was inclusive of both Imd and Toll targets, along with other genes more equally induced across infection conditions. The distinction between the Gram-negative response versus the Gram-positive/fungal response lay in the amplitude of transcriptional upregulation among Imd and Toll targets induced by each type of infection.

The canonical/stereotypical response to Gram-negative versus Gram-positive/fungal pathogens has often been extrapolated, implicitly or explicitly, to mosquitoes (17–22). There is, however, despite early transcriptional analyses (23), a lack of data to either definitively support or disprove this inference. Many components of the Imd and Toll pathways responsible for signal transduction in Drosophila are conserved (24–26), but the transcriptional targets validated in Drosophila are not. Neither have the activations of the pathways been tied to recognition of specific pathogen-associated molecular patterns (PAMPs, *e.g.*, DAP, Lys, β-glucan), host damage, etc. Upregulation of Imd pathway-related genes in mosquitoes (*e.g.*, *REL2*), and/or reduced survival following RNAi-mediated silencing of the Imd pathway have been reported following infection with bacteria of both Gram types (27,28), and with fungi (29–31). The mosquito Toll pathway, in functional studies, has mainly been studied in the context of fungal infection (32–35), although the induction of putative Toll targets (*e.g.*, *cact*, *serpin 27A*) has also been documented following Gram-negative and Gram-positive challenge (28) and systemic *Wolbachia* infection (36,37). The response of putative Imd targets to Gram-positive/fungal infection or of putative Toll targets to Gram-negative infection in no way contradicts the canonical model of transcriptional immune control, as the same phenomenon is documented in *D. melanogaster* (16,38,39). A gene which is heavily targeted by Toll may receive lesser, but still measurable, input from Imd (or from other signaling pathways) and vice versa. What remains uncertain is the extent to which recognition of different types of pathogens stimulates the upregulation of distinct cohorts of genes, because of differential pathway activation.

To better understand the transcriptional response to different types of infection in mosquitoes, we used RNAseq to evaluate transcriptomic changes following challenge with (a) a panel of bacteria, including both Gram-negative and Gram-positive pathogens (b) yeasts and filamentous fungi, and (c) heat-killed doses of a Gram-negative bacterium, a Gram-positive bacterium, and a yeast. To maximize the utility of the resulting dataset, and the applicability of any conclusion to a broad range of mosquito vectors, we performed this experiment using two species: *Aedes aegypti*, of the *culicine* lineage, and *Anopheles gambiae* (*s.l.*), of the *anopheline* lineage. The objectives of this project were to characterize the transcriptional response of each mosquito in terms of (a) *similarities* across pathogens (*i.e.*, is there a core pan-microbial infection response?) (b) *differences* between pathogens (*i.e.*, do responses vary for bacteria versus fungi, Gram-negative versus Gram-positive, low- versus high-virulence pathogens?) and (c) cross-species comparison (*i.e.*, are responses conserved between *Ae. aegypti* and *An. gambiae*? Mosquitoes and fruit flies?).

## Results

### An RNAseq approach to dissect the response of *Aedes aegypti* and *Anopheles gambiae* to infection

To characterize mosquitoes’ transcriptome-level response to systemic infection with bacterial and fungal pathogens, we performed RNAseq on pooled female whole bodies of *Ae. aegypti* of the Liverpool strain and *An. gambiae* (*s.l.*) of the G3 strain. Conditions included unchallenged (UC), mock-wounded (Mock), and systemically-challenged by injection with 3,000 colony-forming units (CFU) of live or heat-killed (HK) pathogen. Live challenges included the Gram-negative bacteria *Escherichia coli* (*Ec*), *Erwinia carotovora carotovora 15* (*Ecc15*), *Serratia marcescens* type strain (*Sm*), and *Providencia rettgeri* (*Pr*); the Gram-positive bacteria *Staphylococcus aureus* (*Sa*), *Micrococcus luteus* (*Ml*), and *Enterococcus faecalis* (*Ef*); and the yeasts *Saccharomyces cerevisiae* (*Sc*) and *Candida albicans* (*Ca*). Microbes were chosen for their range of virulence in *Ae. aegypti*, *An. gambiae*, and *D. melanogaster* (Fig 1A) as measured by survival after systemic challenge (see Fig 1 S1A for mosquitoes, Troha *et al.* for Drosophila (16)). We observed that the mortality incurred by each pathogen was closely correlated in the two mosquito species (R^2^ = 0.95), indicating similar virulence, but that the mortalities associated with some infections in *D. melanogaster* were substantially higher (*Sa*) and lower (*Sm*) than their counterparts in mosquitoes (Fig 1 S1B). Mock wounding and live challenges were assayed at 12 hours and, where survival was sufficiently high, 36 hours post-challenge. HK challenges consisted of the equivalent of 3,000 CFUs of *Ec* or *Ef* (*Ec*HK, *Ef*HK), and were assayed only at 12 hours post-challenge.

**Figure 1:**
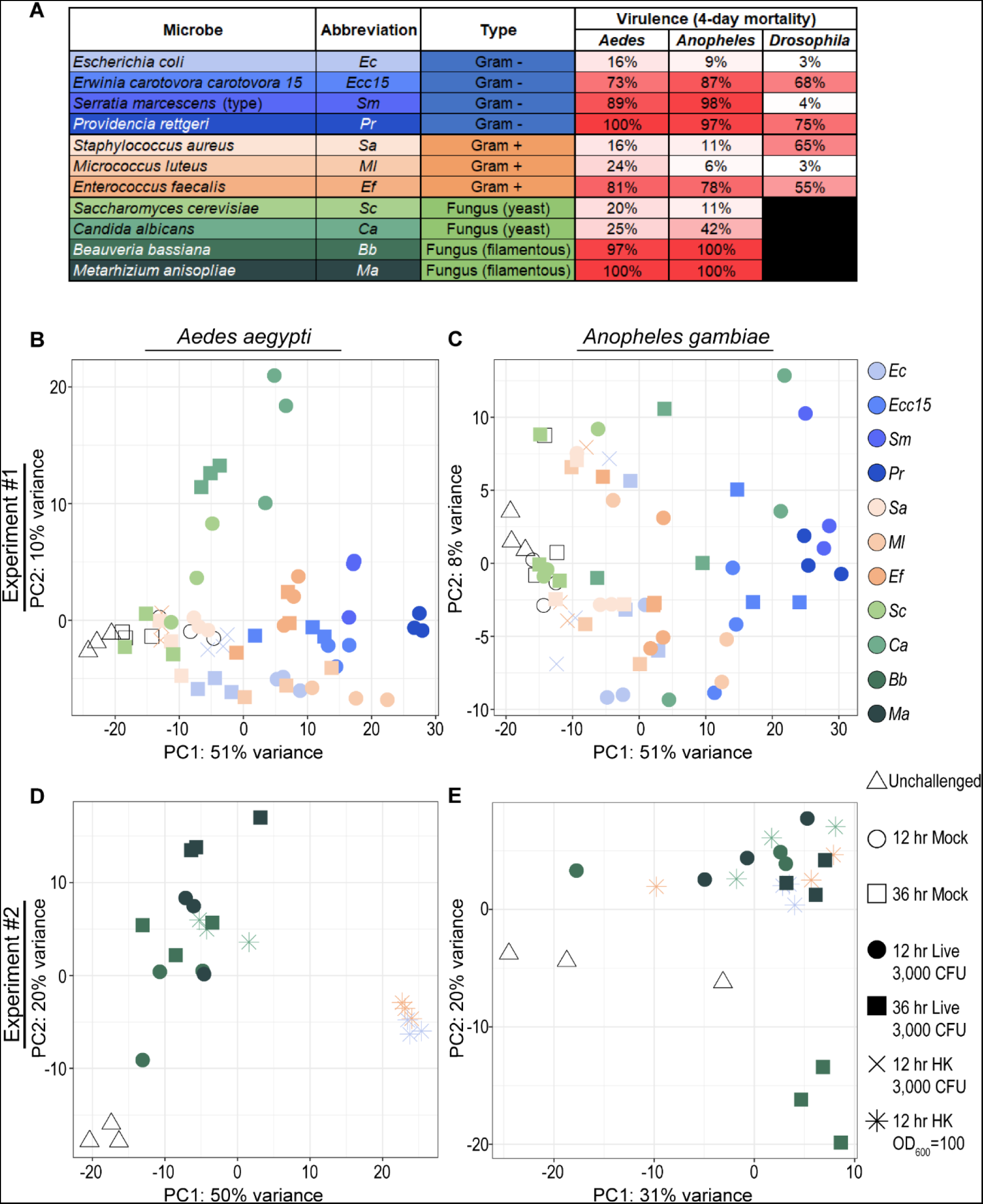
Systemic challenge with pathogens of varying virulence in *Aedes aegypti* and *Anopheles gambiae* (*s.l.*) reveals transcriptional signatures of bacterial and fungal infection. *Aedes aegypti* and *Anopheles gambiae* (*s.l.*) mosquitoes were mock injected or challenged with live and heat-killed (HK) Gram-negative, Gram-positive, and fungal pathogens of varying virulence, here quantified by % mortality at 4-days post challenge (**A**). Data in B-E are the product of two RNAseq experiments. Experiment #1 generated transcriptomes from mosquitoes in the following conditions: unchallenged (UC); mock wounded (Mock, 12- and 36-hour); live-challenged (3,000 colony-forming units, CFUs) with *Escherichia coli* (*Ec*), *Erwinia carotovora carotovora 15* (*Ecc15*), *Serratia marcescens* type strain (*Sm*), *Providencia rettgeri* (*Pr*), *Staphylococcus aureus* (*Sa*), *Micrococcus luteus* (*Ml*), *Enterococcus faecalis* (*Ef*), *Saccharomyces cerevisiae* (*Sc*) and *Candida albicans* (*Ca*) (12- and 36-hour); and challenged with the HK equivalent of 3,000 CFUs of *Ec* and *Ef* (12-hour only). Note that, for *Sm*- and *Pr*-challenged mosquitoes, survival was too low at 36 hours to complete the second timepoint. Experiment #2 generated transcriptomes from mosquitoes in the following conditions: UC; live-challenged (3,000 injected conidia) with *Beauveria bassiana* (*Bb*) and *Metarhizium anisopliae* (*Ma*) (12- and 36-hour); and challenged with OD_600_ = 100 HK *Ec*, *Ef*, and *Ca* (12-hour only). Principal component analyses display the results in *Ae. aegypti* and *An. gambiae*, respectively, of experiments #1 (**B** and **C**) and #2 (**D** and **E**).

A principal component analysis (PCA) (Fig 1 B-C) revealed that the HK-challenged transcriptomes clustered with mock-wounded transcriptomes in both species, indicating that a dosage of *Ec*HK or *Ef*HK equivalent to 3,000 CFUs was not sufficient to generate a distinct PAMP-induced transcriptional signature in either mosquito species. We further noted that, in *Ae. aegypti*, yeast-challenged transcriptomes including *Ca* (12 and 36 hours) and some *Sc* (12 hours) appeared to cluster separately on PC2, which, we hypothesized, might signify a distinct response to fungal infection. To obtain more robust PAMP-induced transcriptomes, and to investigate the potential fungal signature, we performed a second RNAseq experiment with UC mosquitoes, as well as mosquitoes challenged with (a) concentrated *Ec*HK, *Ef*HK, and *Ca*HK (OD_600_ = 100, respectively equivalent to approximately 1.5*10^6^, 3.75*10^6^, and 7.5*10^4^ CFUs of each microbe) at 12 hours and (b) 3,000 conidia of live *Beauveria bassiana* (*Bb*) and *Metarhizium anisopliae* (*Ma*) at 12 and 36 hours (injected, not topically applied, to maximize comparability with other challenges). *Bb* and *Ma*, both filamentous fungi, were selected for their relevance as potential biocontrol agents against mosquitoes (40,41). PCAs were performed on the transcriptomes obtained from the second experiment (Fig 1 D-E). Overall, we observed that, in both *Ae. aegypti* and *An. gambiae*, the intensity of the transcriptional responses elicited by bacterial challenge (as indicated by positioning along PC1) appeared correlated to the virulence of the pathogen (Fig 1 A, B and C). In neither species did we observe clear separation of Gram-negative and Gram-positive challenges, either live (Fig 1B-C) or HK (Fig 1D-E). Finally, in *Ae. aegypti*, but not *An. gambiae*, fungal challenges (live and HK) clustered separately from bacterial challenges (live and HK) (Fig 1 B and D).

### Mosquitoes mount a “core” pan-microbial response to infection

In a previous study from our lab, Troha *et al*. demonstrated that, when systemically challenged, *D. melanogaster* mounts a shared, or “core”, transcriptional response to bacterial pathogens including *Ec*, *Ecc15*, *Sm*, *Pr*, *Sa*, *Ml*, and *Ef*, among others (16). To determine whether *Ae. aegypti* and *An. gambiae* mounted a core response to the microbes in our panel, we first established thresholds to define differentially expressed (DE) genes. For the purposes of this study, a gene is considered DE for a given challenge when it is (a) expressed at a minimum of 1 transcript per million (TPM) in either the UC condition, the given challenged condition, or both and (b) is up or downregulated at least 1.5-fold relative to UC (c) with a DESeq2 *padj* value <0.05.

To examine our data for evidence of a pan-microbial core response to infection, we first analyzed how many genes from each live infection at each timepoint were DE either up or down relative to UC (Fig 2A-B). We noted that, in *Ae. aegypti*, more genes were DE at 12 hours than at 36 hours for most infections, with the most notable exceptions in filamentous fungus-infected mosquitoes (Bb and Ma), where a greater number of DE genes was observed at the later timepoint. In *An. gambiae*, Mock wounding, *Ec*, *Ecc15*, *Sc*, *Bb*, and *Ma* challenge all yielded larger numbers of upregulated genes at 36 hours than at 12 hours. We also observed that the filamentous fungi produced an even higher proportion of DE genes at 36 hours relative to 12 hours post-challenge than in *Ae. aegypti*. These results suggest a difference in the timing of the infection response in the two mosquito species, with *An. gambiae* mounting a slower or more prolonged response than *Ae. aegypti*, which may also be explained by the slightly different rearing conditions used in this study. They also suggest that either filamentous fungal infections proceed more slowly than bacterial infections, or that mosquito hosts are slower to recognize and respond to this type of infection.

**Figure 2:**
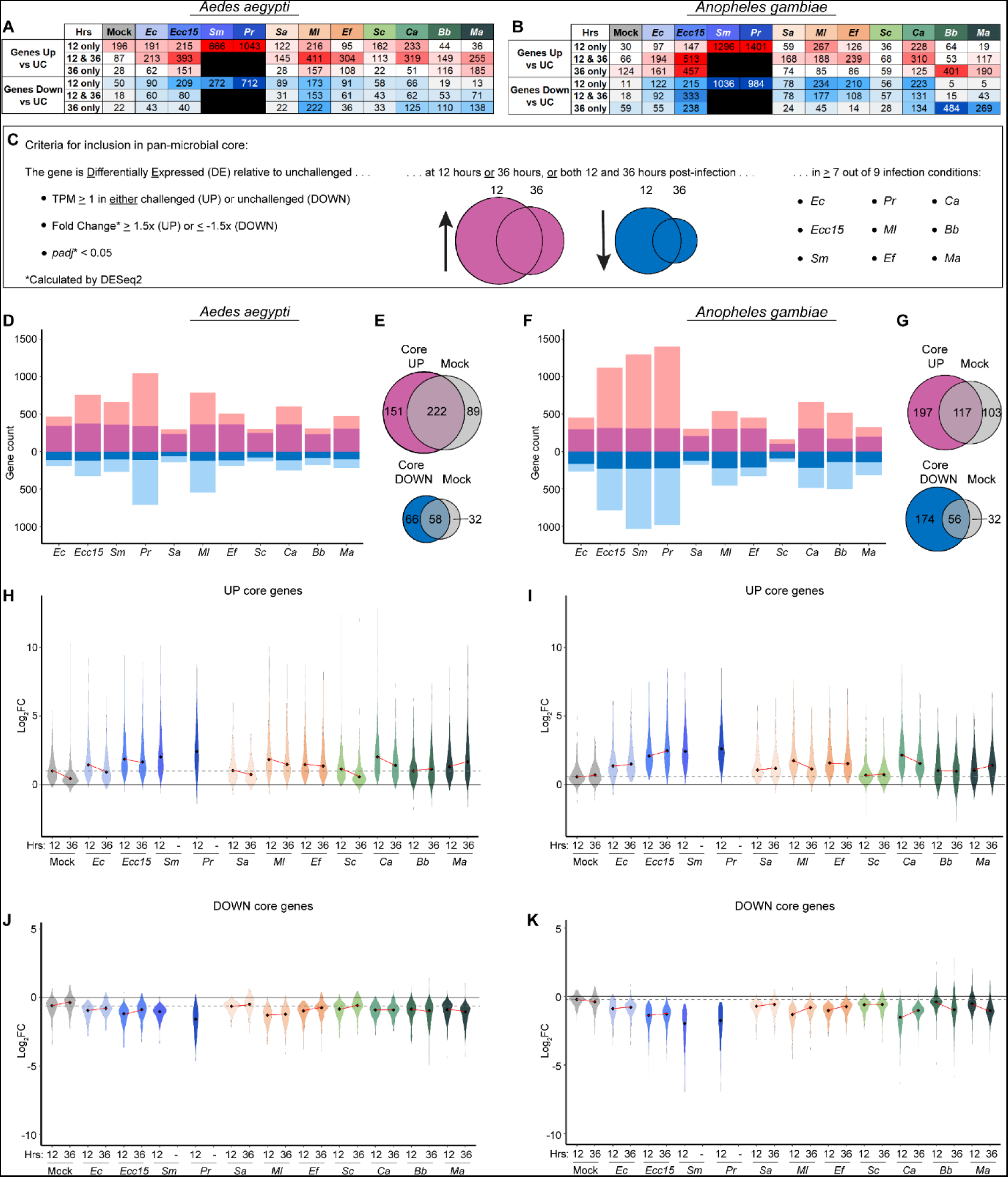
*Aedes aegypti* and *Anopheles gambiae* mount a pan-microbial core transcriptional response to infection with diverse pathogens that is distinct from the wounding response. The number of differentially expressed (DE) genes (DESeq2 fold change ≥ 1.5, *padj* < 0.05) was identified for mock-wounded (Mock) and live-infected *Aedes aegypti* (**A**) and *Anopheles gambiae* (*s.l.*) (**B**) at 12 and 36 hours post-challenge. Live challenges included *Escherichia coli* (*Ec*), *Erwinia carotovora carotovora 15* (*Ecc15*), *Serratia marcescens* type strain (*Sm*), *Providencia rettgeri* (*Pr*), *Staphylococcus aureus* (*Sa*), *Micrococcus luteus* (*Ml*), *Enterococcus faecalis* (*Ef*), *Saccharomyces cerevisiae* (*Sc*), *Candida albicans* (*Ca*), *Beauveria bassiana* (*Bb*), and *Metarhizium anisopliae* (*Ma*). Criteria for a gene’s inclusion in the pan-microbial core are detailed in panel **C**. Bar plots display the number of genes DE in response to live infection (inclusive of both timepoints) in *Ae. aegypti* (**D**) and *An. gambiae* (**F**), including genes in the UP or DOWN pan-microbial cores (dark colors), and those that did not meet the criteria for inclusion (light colors). Venn diagrams display the extent of overlap between the UP cores and DOWN cores (**E** and **G**) for each species with the genes that were DE after mock wounding (inclusive of both timepoints). The log_2_ values of the fold change (relative to unchallenged, calculated by DESeq2) for each gene in the UP (**H** and **I**) and DOWN (**J** and **K**) cores for Mock and live infection conditions at both 12 and 36 hours post-challenge are displayed in violin plots. Black diamonds mark the median value for each condition. The gray dotted line indicates the level of the median log_2_ fold change for the Mock condition at 12 hours post-challenge. The red lines show the trajectory of change in median values between 12 and 36 hours for each condition.

For most infections, a large proportion of DE genes were affected at both timepoints, indicating sustained transcriptional effects (Fig 2A-B, S1). Overall, large numbers of genes DE at 36 hours post-challenge relative to Mock indicate that the transcriptional effects of most infections were ongoing at this timepoint. Accordingly, to more inclusively characterize the core response to infection, we included in our analysis genes that were DE at either timepoint. When all DE genes from the live challenges, inclusive of both timepoints, were compared to the DE genes from the Mock treatment, likewise inclusive of both timepoints (Fig 2 S2), we observed that the *Sa* and *Sc* challenges in both mosquito species were only marginally different from the Mock. This result corroborated the clustering of the corresponding data points previously observed in PCAs (Fig 1B-C). We concluded that, for these microbes, 3,000 CFUs were insufficient to produce a robust infection in *Ae. aegypti* and *An. gambiae*, and we consequently excluded *Sa* and *Sc* from our pan-microbial core analysis. Of the nine remaining infectious challenges we set a minimum threshold of seven. This less restrictive cutoff was set to avoid overlooking genes that failed to meet the DE threshold in one or two infections due to infection-specific dynamics (*e.g.*, early clearance of the pathogen, bacterial suppression of the canonical response, slower kinetics of infection). Therefore, the final criteria for a gene to be considered a part of the pan-microbial core (either up or down) were that it was DE relative to UC, at one or both timepoints (12 or 36 hours post-challenge), for at least seven out of nine (*Ec*, *Ecc15*, *Sm*, *Pr*, *Ml*, *Ef*, *Ca*, *Bb*, and *Ma*) infectious challenges (Fig 2C). These criteria yielded a pan-microbial core response of 373 upregulated and 124 downregulated genes in *Ae. aegypti*, and 314 upregulated and 230 downregulated genes in *An. gambiae*. For the complete list of all pan-microbial core genes, see Table S1.

We observed that, with the exception of *Sc* infection in *An. gambiae*, which had a much smaller overall number of DE genes as compared to all other treatments, most of the genes in the pan-microbial core were DE in all live infection conditions (Fig 2D and F), indicating that the criteria for inclusion were sufficiently stringent to capture a true shared transcriptional response across infection conditions. In both species, the upregulated core and downregulated core overlapped with genes upregulated and downregulated in the Mock condition (inclusive of both timepoints) (Fig 2E and G), raising the possibility that a large portion of the core response we characterized might be a reaction to wounding rather than infection. However, when the amplitude of the fold-change (calculated by DESeq2, relative to UC) of genes in the upregulated cores (Figs 2H-I) and downregulated cores (Figs 2J-K) was compared across conditions/timepoints, we found that the median change in most infection conditions was greater than in Mock at either timepoint, confirming that infection played a role, over and above wounding, in altering the expression of core genes. This higher amplitude of change in live infection conditions relative to Mock is also illustrated in heatmaps comparing fold change (calculated by DESeq2, relative to UC) across conditions (Fig 2 S3). It should be noted that, since the Mock wounding was performed in non-sterile conditions, it is impossible to say what portion of the Mock response is attributable to wounding and what resulted from the introduction of environmental microbes from the cuticle surface to the hemocoel. Altogether our data demonstrate that *Ae. aegypti* and *An. gambiae* each mount a core response to infection, and that this response is distinct from the wounding response.

### The upregulated pan-microbial core is conserved between *Aedes aegypti* and *Anopheles gambiae*

To assess the functional signature of the pan-microbial cores in *Ae. aegypti* and *An. gambiae*, we performed an enrichment analysis to identify overrepresentation in the upregulated cores of gene ontology (GO) terms, Interpro domains, and defense-related categories. Defense-related annotations were taken from a list previously published in Hixson et al. (42) updated in this study (see Table S2). Here, our analysis of the enrichment of defense-related genes excludes “low confidence” annotations. For the complete analysis, including GO terms, Interpro IDs, immune gene categories, associated gene IDs, and *padj* values, see Table S3. Selected categories are shown in Fig 3A. We found that, in both species, most enriched categories could be divided into broad functions, including defense, protein folding and secretion, macromolecule biosynthesis/catabolism, and energy metabolism.

**Figure 3:**
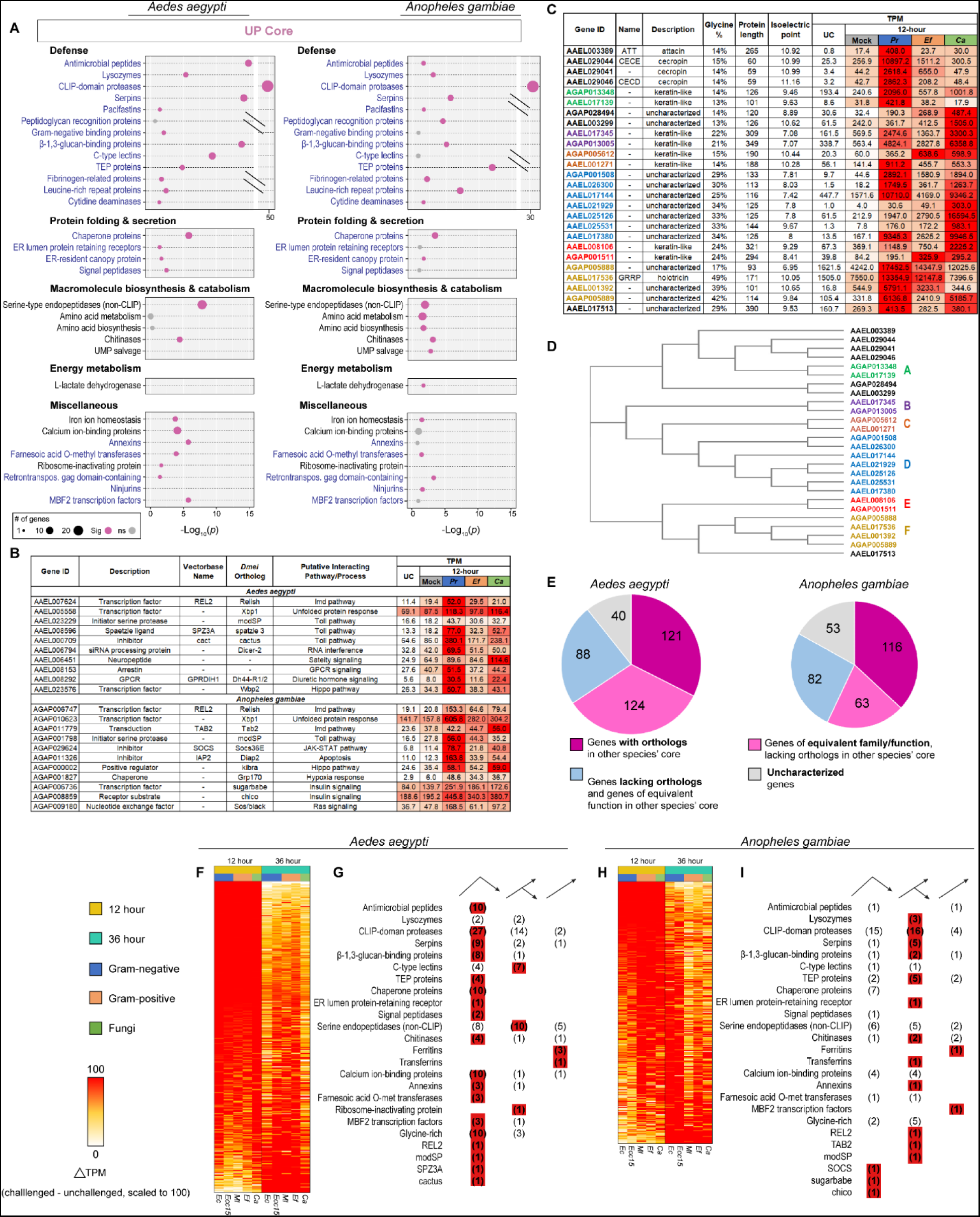
The upregulated core response to infection is well conserved between *Aedes aegypti* and *Anopheles gambiae*. Bubble plots (**A**) display categories of genes enriched in the upregulated pan-microbial cores of *Ae. aegypti* and/or *An. gambiae* (*s.l.*). Black text indicates category is from TopGO. Blue text indicates custom category (either from immune gene list, or defined by the presence of InterPro domain(s)). The size of the bubble is proportional to the number of genes from the given category in the upregulated core. The placement of the bubble along the x axis corresponds to the statistical significance of the enrichment (Fisher’s exact test). Gray bubbles indicate *p* value ≥ 0.05 (not significant enrichment). (**B**) Table of selected genes in the upregulated cores of *Ae. aegypti* and *An. gambiae*. (**C**) Table of uncharacterized glycine-rich (>13%) genes and known glycine-rich antimicrobial peptides in the upregulated cores of *Ae. aegypti* and *An. gambiae*. (**D**) A dendrogram showing sequence similarity between glycine-rich genes from the *Ae. aegypti* and *An. gambiae* cores, generated by Clustal Omega. Genes are grouped by sequence similarity across species into “types” A, B, C, D, E, and F. (**E**) Pie charts describing the orthology and functional similarities shared by the *Ae. aegypti* and *An. gambiae* upregulated cores. Heatmaps of change in expression (challenged TPM minus unchallenged TPM) of core upregulated genes in *Ae. aegypti* (**F**), and *An. gambiae* (**H**) challenged with *Escherichia coli* (*Ec*), *Erwinia carotovora carotovora 15* (*Ecc15*), *Micrococcus luteus* (*Ml*), *Enterococcus faecalis* (*Ef*), and *Candida albicans* (*Ca*), scaled by infection, grouped by timepoint. Tallies of genes in the descending (peaked before 36 hours for at least 4/5 of conditions), ascending (peaked after 36 hours for at least 4/5 conditions), and intermediate (met neither of the previous criteria) cohorts in *Ae. aegypti* (**G**) and *An. gambiae* (**I**).

Defensive functions were well represented in both species’ upregulated cores. We observed an enrichment of antimicrobial peptides (AMPs) and lysozymes. The AMP enrichment was more pronounced in *Ae. aegypti*, comprising ten genes (*ATT*, *CECD*, *CECE*, *CECN*, an unnamed cecropin, *DEFA*, *DEFC*, *DEFD*, *GAM1*, and *GRRP*), compared to two (*CEC1* and *DEF1*) in *An. gambiae*. Lysozyme enrichment was more congruent between the two species, with four genes in *Ae. aegypti* (*LYSC11*, *LYSC7B*, *LYSC4*, and one unnamed gene) and three in *An. gambiae* (*LYSC1*, *LYSC2*, and *LYSC4*). Both species displayed highly significant enrichments of CLIP-domain serine proteases and their proteolytically inactive homologs (forty-three in *Ae. aegypti*, thirty-five in *An. gambiae*) and serpins (twelve in *Ae. aegypti*, six in *An. gambiae*). Both cores contained a single pacifastin serine protease inhibitor (orthologous across the two species); pacifastins have previously been proposed to play a role in arthropod immunity (43,44). Both species’ upregulated cores included pattern-recognition and opsonizing proteins, including amidase PGRPs (*PGRPS1* in both species, and *PGRPLB*, *PGRPS2*, and *PGRPS3* in *An. gambiae*), GNBPs (a single unnamed GNBP in *An. gambiae*, *GNBPA1* and *GNBPB1* in *Ae. aegypti*), putative β-1,3-glucan-binding proteins (nine in *Ae. aegypti*, four in *An. gambiae*), C-type lectins (eleven in *Ae. aegypti*, two in *An. gambiae*), thioester-containing proteins (four in *Ae. aegypti* and nine in *An. gambiae*, including *TEP1*), fibrinogen-related proteins (four in *Ae. aegypti*, five in *An. gambiae*), and leucine-rich immune proteins (four in *Ae. aegypti*, eight in *An. gambiae*). We also observed an orthologous pair of cytidine deaminases (one in each species) which may play a role in innate immunity against viruses. Interestingly, we observed that in *An. gambiae*, pro-phenoloxidases were enriched in the downregulated core (four genes: *PPO2*, *PPO5*, *PPO6*, and *PPO9*) (Fig 3 S1A). A single CLIP-domain serine protease (*CLIPA9*) also appeared in the *An. gambiae* downregulated core (Fig 3 S1B).

In both species’ upregulated cores, we saw the enrichment of genes associated with protein folding, secretion, and the endoplasmic reticulum (ER). These included heat shock/chaperone proteins (ten in *Ae. aegypti*, seven in *An. gambiae*), an orthologous pair of ER lumen protein-retaining receptors (one in each species), an orthologous pair of ER canopy proteins (one in each species), and signal peptidases (two in *Ae. aegypti*, one in *An. gambiae*).

We saw enrichments of multiple categories of enzymes putatively involved in metabolism in the upregulated cores. These included serine endopeptidases lacking regulatory CLIP domains (twenty-three in *Ae. aegypti*, thirteen in *An. gambiae*). The precise functions of most of these peptidases are uncharacterized, with the exception of *LT1*, which is known for its role in blood meal digestion and is transcribed in the gut in response to ecdysone signaling (45). Transcriptomic data from the *Aegypti Atlas* demonstrate that most of the *Ae. aegypti* non-CLIP serine proteases in the upregulated core (apart from *LT1*) are not expressed in the posterior midgut, before or after blood-feeding, and are therefore unlikely to serve a diet-digestion function. Rather, most were primarily expressed in the thorax, abdomen, and head, suggesting an association with the fat body (42). In *An. gambiae* we also observed an enrichment of enzymes associated with diverse processes related to amino acid biosynthesis/metabolism. There was no significant enrichment of these categories in *Ae. aegypti*, however, we noted the following orthologous pairs shared in both cores: AAEL017029 and AGAP005712 (aromatic amino acid hydroxylases), AAEL000271 and AGAP006670 (gamma-glutamyl hydrolases). Both species’ cores were enriched with chitinases (six in *Ae. aegypti*, five in *An. gambiae*). *An. gambiae* expressed two enzymes involved in uridine salvage (one uridine kinase and one uridine phosphorylase).

With respect to energy metabolism, L-lactate dehydrogenase was upregulated in *An. gambiae,* while, in the downregulated cores, we observed enrichments of mitochondrial transporters and pyruvate metabolizing enzymes in both species, and of NADH dehydrogenases, cytochrome c oxidases, and ATP synthase subunits in *An. gambiae* (Fig 3 S1A). These results suggest the suppression of aerobic respiration and, in *An. gambiae*, an increased reliance on anaerobic respiration.

Among upregulated core genes, we also observed a range of miscellaneous categories, significantly enriched in one or both species, which merit comment. These included genes involved in iron ion homeostasis comprising ferritin subunits (three in *Ae. aegypti*, one in *An. gambiae*) and an orthologous pair of transferrins (one in each species). The upregulation of iron-binding proteins likely serves to sequester iron to limit microbial growth (46). Calcium-binding proteins, including gelsolins, calreticulins, and annexins were also upregulated in both cores (fourteen in *Ae. aegypti*, eight in *An. gambiae*), indicating that calcium signaling plays a role in the infection response. We also observed the upregulation of farnesoic acid O-methyl transferases (three in *Ae. aegypti*, two in *An. gambiae*), which are believed to catalyze the rate-limiting step in the synthesis of methyl farnesoate, an intermediate in the synthesis of juvenile hormone (47,48). *Ae. aegypti* but not *An. gambiae* expressed an rRNA N-glycosylase (*i.e.*, ribosome-inactivating protein). The protein encoded by this gene contains a domain (PF00161), which inactivates eukaryotic ribosomes. While this domain is most commonly found in plant and bacterial toxins (*e.g.*, ricin and shiga toxin), it also is present in the genome of *culicine* mosquitoes, as the legacy of a horizontal gene transfer from a cyanobacterium (49,50). Ribosome-inactivating proteins have been proposed to serve an antiparasitic immune function in *Ae. aegypti* (51). The core upregulated responses of both species contained three retrotransposon gag domain-containing proteins (one in *Ae. aegypti*, two in *An. gambiae*). We speculate that the presence of virus-derived sequences in the upregulated core indicates a loss of control over the expression of selfish genetic elements during the stress of infection. The *An. gambiae* upregulated core included two ninjurin proteins, which are regulated by injury and stress in *D. melanogaster* (52). Finally, MBF2 transcription factors were present in both species’ upregulated cores (four in *Ae. aegypti*, one in *An. gambiae*).

Overall, enrichment analysis demonstrated that, among upregulated core genes, defensive functions are strongly emphasized. The increased expression of chaperones and ER-associated proteins likely relates to the heightened output of secreted proteins (*e.g.*, AMPs) instigated by the activation of immune signaling pathways in response to infection. The altered expression of metabolic genes may point to changing metabolic priorities which could help the mosquito host tolerate infection.

Further examination of the core upregulated responses revealed additional genes of interest, not captured by enrichment analysis (Fig 3B). *REL2*, the terminal transcription factor of the IMD pathway was present in both cores, suggesting that *REL2* targets itself for transcriptional upregulation upon activation of Imd signaling. Both cores also contained orthologs of the transcription factor *Xbp1*, part of the unfolded protein response in *D. melanogaster* (53), a further sign of ER stress in the infection response. In the *Ae. aegypti* core, we observed *SPZ3A* (a putative Toll pathway cytokine) and the Toll inhibitor *cact* (possibly transcribed downstream of Toll activation as part of a self-modulating feedback loop). Additional genes of interest included *Dicer-2* (suggesting transcriptional crosstalk between immune signaling pathways and the siRNA pathway), a putative neuropeptide with a conserved gastrin/cholecystokinin site (possibly mediating an anorexic response to infection), an arrestin, *GPRDIH1* (a GPCR involved in diuretic hormone signaling), and an ortholog of the *D. melanogaster* transcription factor *Wbp2* (a partner of *yorkie* in the hippo pathway). In the *An. gambiae* core, we found the IMD component *TAB2*, the repressor *SOCS* (possibly transcribed downstream of the *JAK-STAT* pathway as part of a self-modulating feedback loop), *IAP2* (an inhibitor of apoptosis), an ortholog of *kibra* (a positive regulator of the hippo pathway in *D. melanogaster*), an ortholog of the *D. melanogaster* chaperone protein *Grp170* (expressed in response to hypoxic stress), orthologs of the *D. melanogaster* genes *sugarbabe* and *chico* (components of the insulin signaling pathway), and an ortholog of the *D. melanogaster* gene *Son of sevenless* (a nucleotide exchange factor involved in the activation of Ras).

In both mosquitoes’ upregulated cores, we noted a common theme of proteins with high glycine content. Overrepresentation of glycine residues has been identified as a common characteristic in certain insect AMPs, including attacins and diptericins(54). Several of the known AMPs in the *Ae. aegypti* core response were glycine-rich, with contents ranging from 14-49%. These observations prompted us to take a census of glycine-rich peptides in the upregulated core pan-microbial response. We selected all genes (a) with glycine content ≥ 13%, (b) with a signal peptide sequence, and (c) lacking clear functional annotation. These criteria yielded thirteen genes in *Ae. aegypti* and eight in *An. gambiae*. The known glycine-rich AMPs in the core (*ATT*, *CECE*, AAEL29041 (an unnamed cecropin), *CECD*, and *GRRP*) were included for comparison (Fig 3C). We noted that several of the uncharacterized proteins in our census were predicted to possess high isoelectric points (>10), comparable to the known AMPs in our analysis. Positive charge is another characteristic associated with AMP function (55). We used Clustal Omega (56,57) to group the polypeptide sequences of all twenty-six genes by similarity (Fig 3D). We then performed pairwise comparisons of neighboring genes by polypeptide alignment. In this manner, we sorted the majority of the uncharacterized glycine-rich proteins in our census into pairs or larger groups, containing one or more genes from each species, with highly similar amino acid sequences. For ease of communication, we have designated these groups as “types” A, B, C, D, E, and F (order is arbitrary, see Fig 3 S2A-F for alignments). Apparent orthologs to some of these groups were also identified in *D. melanogaster* and *Culex quinquefasciatus* (Fig 3 S2J). Two of the glycine-rich types we identified were of special interest. Type D includes a single *An. gambiae* gene, and six *Ae. aegypti* paralogs. The six *Ae. aegypti* Type D glycine-rich paralogs are located together on chromosome 3 (Fig 3 S2G). We were also intrigued to discover that the *Ae. aegypti* AMP *GRRP* shares apparent sequence similarity with its neighboring gene on chromosome 2 (AAEL001392, Fig 3 S2H) which, in turn appeared highly similar to the two *An. gambiae* genes, AGAP005888 and AGAP005889, which neighbor each other on chromosome 2 (Fig 3 S2I). We propose that these four genes are homologous and hereafter in this study collectively designate them “Type F glycine-rich proteins”. Overall, we found that the core responses of *Ae. aegypti* and *An. gambiae* featured an abundance of homologous uncharacterized glycine-rich peptides, some of which exhibit properties (predicted isoelectric point) or sequences similar to those of known AMPs.

In assessing the upregulated cores in *Ae. aegypti* and *An. gambiae*, we found the composition of the two cores remarkably similar, suggesting a conserved infection response. To confirm this conservation, we examined the two upregulated cores for orthology, using the VectorBase orthology search function. We found that 121 genes in the *Ae. aegypti* upregulated core (32.5%) possessed at least one ortholog in the *An. gambiae* upregulated core, and another 124 genes (33.2%) either belonged to the same family (*e.g.*, CLIP-domain serine proteases, C-type lectins) or shared an equivalent molecular function (*e.g.*, chitinases, peroxidases) with a member of the *An. gambiae* upregulated core (Fig 3E). Only 88 genes (23.6%) with a clearly characterizable function lacked a counterpart in *An. gambiae*. In the *An. gambiae* upregulated core, 116 genes (36.9%) shared orthology with an *Ae. aegypti* core gene, 63 (20.1%) shared at least one family/functional counterpart, and only 53 (16.9%) characterizable genes lacked any counterpart in the *Ae. aegypti* upregulated core. Overall, we found that *Ae. aegypti* and *An. gambiae* mount core responses to infection which are conserved with respect to both function and orthology.

### Timing of the core response

Having defined the functions of the upregulated cores, we next sought to characterize the timing of their expression. Since the cores were defined using aggregates of genes upregulated at 12 and 36 hours post-infection (PI), it was necessary to disaggregate them in order to describe their timing. For this analysis, we excluded low-response infections (*Sa* and *Sc*) as well as single timepoint infections (*Sm* and *Pr*) leaving *Ec*, *Ecc15*, *Ml*, *Ef*, *Ca*, *Bb*, and *Ma*. We first scaled the expression of each core gene within each infection according to whether it was higher expressed at 12 hours or 36 hours. The timepoint with the higher expression was set to 100, with the lesser timepoint expressed as a percentage of 100. Heatmaps of the resulting patterns of expression demonstrated that most core genes were higher expressed at 12 hours for *Ec*, *Ecc15*, *Ml*, *Ef*, and *Ca* infections, but for *Bb* and *Ma* infections more genes were higher expressed at 36 hours (*Aedes*: Fig 3 S3 *Anopheles*: Fig 3 S4). Consequently, we divided the data and performed one analysis for the bacteria and *Ca*, and a separate analysis for the filamentous fungi. We next divided the core genes into *descending* genes (higher expressed at 12 hours in at least four out of five bacterial/*Ca* infections; two out of two for filamentous fungi) *ascending* genes (higher expressed at 36 hours in at least four out of five bacterial/Ca infections; two out of two for filamentous fungi), and *intermediate* genes (higher expressed at 12 or 36 hours for no more than three bacterial/*Ca* infections; oppositely oriented in *Bb* and *Ma* for filamentous fungi). For the filamentous fungi, genes which were not significantly regulated at least 1.5-fold by both *Bb* and *Ma* infection at a minimum of one timepoint were excluded from the analysis. We assumed that the descending genes reached peak expression before 36 hours, the ascending genes reached peak expression sometime after 36 hours, and the intermediate genes peaked near 36 hours. We then quantified how the functional categories and notable genes identified in our previous analyses were distributed with respect to timing. The full results of our analysis for bacteria/yeast and filamentous fungal infections can be found in Fig 3 S3 (*Ae. aegypti*) and Fig 3 S4 (*An. gambiae*). Figure 3F-I displays a condensed summary of the results of the bacteria/*Ca* analysis of the upregulated core. Overall, we found that most functional categories we identified in the upregulated core belonged to the descending set in *Ae. aegypti*, and to the intermediate set in *An. gambiae*, indicating that, under the conditions we employed, most of the core response in *Ae. aegypti* peaked before 36 hours, while the *An. gambiae* response was either induced more gradually or upregulated in a more sustained manner. It is noteworthy that the iron-binding categories (ferritins and transferrins) were mainly expressed in the ascending set for both species, indicating that, in contrast to immune effectors, iron sequestration is a lagging or prolonged component of the infection response. As we observed in our analysis of DE genes, the transcriptional response to the filamentous fungi was also slower in both species than the response to bacteria/*Ca*, likely either reflecting slower pathogen growth or slower recognition of fungal infection.

### *Aedes aegypti* and *Anopheles gambiae* do not discriminate between Gram-negative and Gram-positive bacterial infection

In *D. melanogaster*, differential activation of the Imd and Toll pathways, respectively, mediate differential responses to Gram-negative and Gram-positive bacteria. To determine whether the immune response in *Ae. aegypti* and *An. gambiae* also discriminates by Gram type, we probed the transcriptomes of *Ec*-, *Ecc15*-, *Sm*-, and *Pr*-infected mosquitoes for differences from the transcriptomes of *Sa-*, *Ef-*, and *Ml*-infected mosquitoes, using equivalent transcriptomes from *D. melanogaster* (16) as a positive test-case for comparison. We first compared all transcriptomes for the selected microbes, genome-wide, by PCA. In *D. melanogaster*, the transcriptomes of Gram-negative-infected flies were completely separated from the transcriptomes of Gram-positive-infected flies on PC2, accounting for approximately 20% of the variance in the comparison (Fig 4A). The PCAs of *Ae. aegypti* (Fig 4B) and *An. gambiae* (Fig 4C) transcriptomes were markedly similar to one another. Neither exhibited clear separation of Gram-negative-infected transcriptomes from Gram-positive-infected transcriptomes on either PC1 or PC2. While Gram-negative-infected transcriptomes were, on average, farther removed from UC transcriptomes on PC1, this could be an artifact of confounding, as the Gram-negative pathogens were also, on average, more virulent than their Gram-positive counterparts (Fig 1A). To eliminate the signature of virulence from the infection, we performed a separate RNAseq experiment comparing the effects of inoculation with large doses (OD_600_ = 100) of *Ec*HK and *Ef*HK. When the resulting transcriptomes were compared by PCA, we found no separation of the two treatments on either PC1 or PC2.

**Figure 4:**
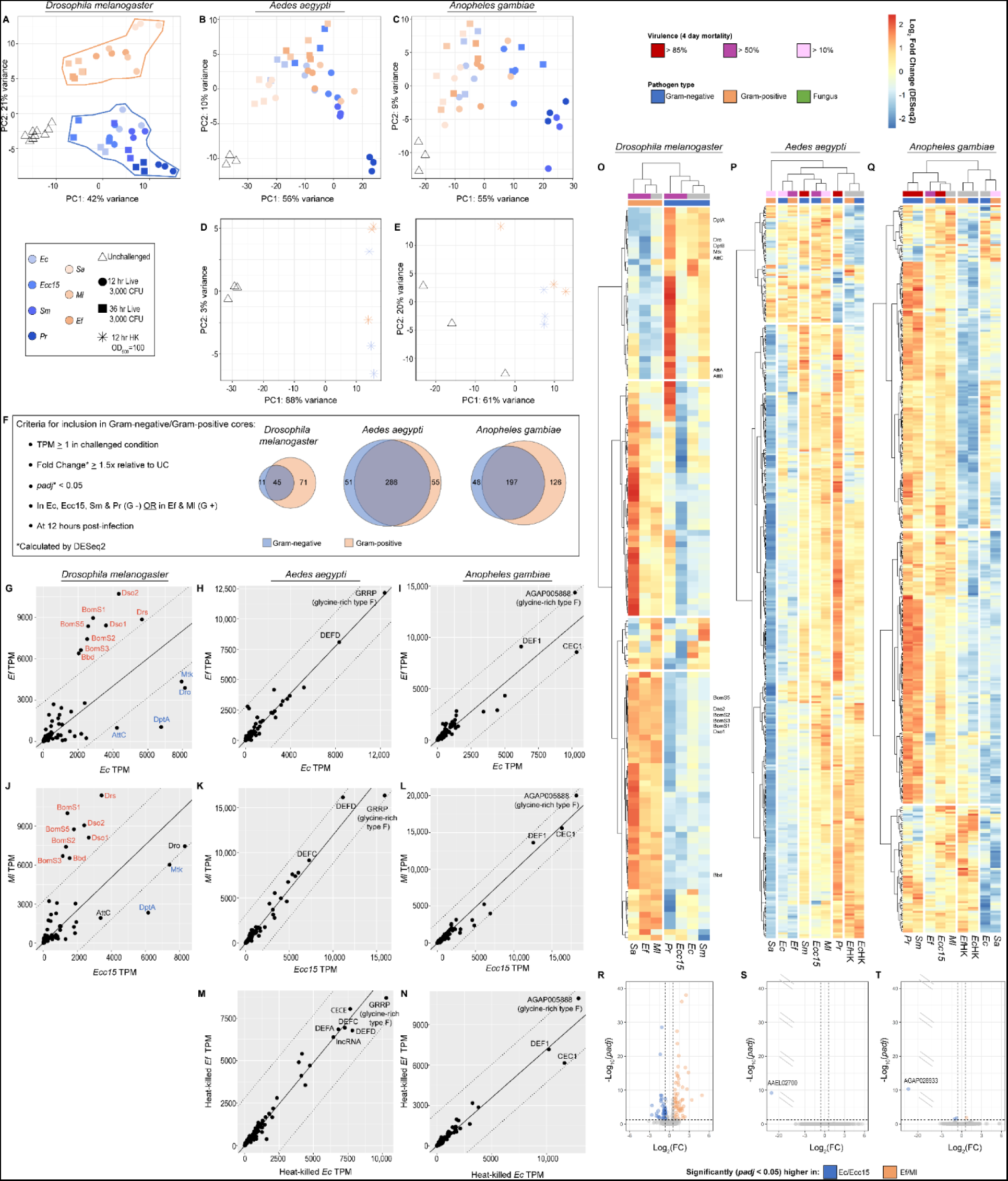
*Aedes aegypti* and *Anopheles gambiae* do not discriminate transcriptionally between Gram-negative and Gram-positive infections. Principal component analyses of transcriptomes from Gram-negative- and Gram-positive-challenged *Drosophila melanogaster* (**A**), *Ae. aegypti* (**B**), and *An. gambiae* (*s.l.*) (**C**). Challenges included *Escherichia coli* (*Ec*), *Erwinia carotovora carotovora 15* (*Ecc15*), *Serratia marcescens* type strain (*Sm*), *Providencia rettgeri* (*Pr*), *Staphylococcus aureus* (*Sa*), *Micrococcus luteus* (*Ml*), *Enterococcus faecalis* (*Ef*), *Saccharomyces cerevisiae* (*Sc*) and *Candida albicans* (*Ca*). Principal component analyses of transcriptomes from *Ae. aegypti* (**D**) and *An. gambiae* (**E**) challenged with concentrated (OD_600_ = 100) heat-killed *Ec* and heat-killed *Ef*. (**F**) Criteria for inclusion in the Gram-negative and Gram-positive cores, with Venn diagrams showing the size and relationship between cores in *D. melanogaster*, *Ae. aegypti*, and *An. gambiae*. Plots depicting the expression (in transcripts per million) of genes in the Gram-negative and Gram-positive core genes in *D. melanogaster* (**G**, **J**), *Ae. aegypti* (**H**, **K**, **M**) and *An. gambiae* (**I**, **L**, **N**) juxtaposing the response to *Ec* versus *Ef* (**G**, **H**, **I**), *Ecc15* versus *Ml* (**J**, **K**, **L**), and heat-killed *Ec* versus heat-killed *Ef* (**M**, **N**); all data from 12-hour timepoint. Linear regressions (solid lines) are plotted through the origin. Dotted lines delineate an arbitrary perpendicular geometric distance of 2000 from the trendline. Clustering analyses of Gram-negative and Gram-positive core genes in *D. melanogaster* (**O**), *Ae. aegypti* (**P**), and *An. gambiae* (**Q**) comparing log_2_FC (calculated by DESeq2) in insects challenged with *Ec*, *Ecc15*, *Sm*, *Pr*, *Sa*, *Ef*, and *Ml*; all data from 12-hour timepoint. Volcano plots depicting genes that are differentially expressed (DESeq2 fold-change ≥ 1.5, *padj* < 0.05) in a comparison of *Ec*- and *Ecc15*-challenged versus *Ef-* and *Ml*-challenged *D. melanogaster* (**R**), *Ae. aegypti* (**S**), and *An. gambiae* (**T**); all data from 12-hour timepoint.

Having failed to identify a Gram type-specific signature in mosquito transcriptomes genome-wide, we next sought to define a smaller core of candidate genes to test for a more subtle Gram type-specific response. From each insect species we selected all genes expressed ≥ 1 TPM that were statistically significantly (DESeq2 *padj* < 0.05) upregulated at least 1.5-fold at 12 hours PI in *Ec*-, *Ecc15*-, *Sm*-, and *Pr*-challenged insects (Gram-negative core) or in *Ef*- and *Ml*-challenged insects (Gram-positive core). *Sa* challenge was excluded, as in the pan-microbial core, as the low-amplitude of the response to this microbe at the 3,000 CFU dose rendered a criterion of differential expression in response to *Sa* overly restrictive for our purposes. Our criteria yielded Gram-negative cores of 56, 337, and 245 genes; and Gram-positive cores of 116, 341, and 323 genes, respectively, in *D. melanogaster*, *Ae. aegypti*, and *An. gambiae*. In all three species, the Gram-negative and Gram-positive cores overlapped substantially (Fig 4F) yielding combined test sets of 127 genes (*D. melanogaster*), 392 genes (*Ae. aegypti*), and 371 genes (*An. gambiae*).

To view the expression of Gram-negative/Gram-positive core genes in comparable infection conditions, we plotted their expression (in TPM) in *Ec*-infected versus *Ef*-infected mosquitoes (Fig 4 G-I) and in *Ecc15*-infected versus *Ml*-infected mosquitoes (Fig 4 J-L) with linear regressions plotted through the origin to represent the overall trend of the data. The pairs of conditions to compare were selected on the basis of similarity with respect to the numbers of genes upregulated at 12 hours PI in *Ae. aegypti* and *An. gambiae* (see Fig 2A-B). In *D. melanogaster*, both plots (Fig 4 G and J) contained two sets of genes at a large distance from the trendline: *Bbd*, *BomS1*, *BomS2*, *BomS3*, *BomS5*, *Drs*, *Dso1*, and *Dso2* (above the trendline, *i.e.*, more responsive to Gram-positive challenge) and *AttC*, *DptA*, *Dro*, and *Mtk* (below the trendline, *i.e.*, more responsive to Gram-negative challenge). The former cohort are documented targets of the Toll pathway (58–60), while the latter four genes are targeted by the Imd pathway (61,62). We calculated the perpendicular geometric distance of each data point from the trendline (see Fig 4 S1 for method) and found that these outlying clusters of Imd and Toll target genes fell a distance >2000 from the trendline. In *Ae. aegypti* and *An. gambiae*, by contrast, the expression of all high-expressed core genes was closely correlated between *Ef* and *Ec* (Fig 4H and I), and between *Ml* and *Ecc15* (Fig 4K and L). Correlation of the expression of core genes in *Ec*HK- versus *Ef*HK-challenged mosquitoes (Fig 4M-N) likewise yielded no clear dissimilarities between the responses to Gram-negative versus Gram-positive PAMPs. All data points in these plots lay within a perpendicular geometric distance from the trendline of less than 2000. While 2000 as a cutoff for differential regulation in the compared conditions is somewhat arbitrary, all genes beyond this threshold, in Drosophila comparisons, were found to be well-documented Imd and Toll pathway targets. For mosquitoes, we believe a distance of 2000 is a conservative threshold to adopt, given the higher overall amplitude of expression in *Ae. aegypti* and *An. gambiae* than in *D. melanogaster*. Distance scores for all data points can be found in Table S4. Overall, this analysis reaffirmed the existence of a differential response to Gram-negative versus Gram-positive infection in *D. melanogaster* and failed to demonstrate a similar distinction in the responses of *Ae. aegypti* and *An. gambiae*. On an unrelated note, we were interested to observe that, in both mosquito species, the highest expressed gene in all conditions was a type F glycine-rich protein (*GRRP* in *Ae. aegypti*, AGAP005888 in *An. gambiae*), reinforcing the similarity between these genes which we had noted previously (see Fig 3C-D, S2F).

While high-expressed genes, such as AMPs, are clearly visible in a plot of TPMs, a more modestly expressed Gram type-specific response might be overlooked. Accordingly, for a more egalitarian comparison, we performed a hierarchical clustering analysis on each species’ Gram-negative/Gram-positive core genes to compare log_2_fold change across Gram-negative and Gram-positive conditions. In *D. melanogaster* (Fig 4O) we found that the first hierarchical division was between the Gram-negative and Gram-positive infection conditions. We also observed one cluster of genes, most strongly upregulated by Gram-negative infection conditions, containing the Imd pathway targets *DptA*, *Dro*, *DptB*, *Mtk*, *AttC*, *AttA*, and *AttB*. Another cluster, most strongly upregulated by Gram-positive infection conditions, contained the Toll pathway targets *BomS5*, *Sdo2*, *BomS2*, *BomS3*, *BomS1*, *Dso1*, and *Bbd*. In *Ae. aegypti* (Fig 4P) and *An. gambiae* (Fig 4Q), hierarchical clustering did not appear to be influenced by the Gram type of the infections.

In a final attempt to identify a Gram-negative or Gram-positive-specific infection response, we initially used DESeq2 analysis to directly compare the genome-wide transcriptomes of mosquitoes challenged by *Ec*HK and *Ef*HK (OD_600_ = 100, Fig 4 S2). We discovered a small number of DE genes in *Ae. aegypti* (Fig 4 S2A), and an even smaller number in *An. gambiae* (Fig 4 S2B). While this differential expression might seem to demonstrate a Gram type-specific response, we noted that comparisons of *Ae. aegypti* challenged with Gram-negative versus Gram-negative bacteria (*e.g.*, *Ecc15* versus *Sm*, Fig 4 S2C) or with Gram-positive versus Gram-positive bacteria (*e.g.*, *Ef* versus *Ml*, Fig 4 S2D) demonstrated similar or even greater levels of differential expression, indicating that there is substantial variability in head-to-head comparisons of individual challenges which is not dependent on Gram type. Upon further examination, we found that the genes which were differentially expressed in the comparison of *Ec*HK to *Ef*HK were not consistently regulated in a Gram-type-specific manner (Fig 4 S2E). Overall, we found that head-to-head comparisons of individual challenges by DESeq2 were not dispositive in establishing whether mosquitoes mount a Gram type-specific response to infection. For a more robust comparison, therefore, we aggregated *Ec*- and *Ecc15*-challenged transcriptomes and compared them to *Ef*- and *Ml*-challenged transcriptomes in a single DESeq2 analysis. In *D. melanogaster* (Fig 4R) this analysis identified large quantities of DE genes. In *Ae. aegypti* (Fig 4S) we found a single gene (AAEL027000, an ortholog of *CRIF* in *D. melanogaster*) which was statistically significantly higher expressed in *Ec*- and *Ecc15*-challenged mosquitoes. However, upon further examination, we found this gene was inconsistently expressed across replicates, and showed no Gram type-specific pattern in the broader dataset. In *An. gambiae* (Fig 4T), we identified four statistically significantly DE genes. Of these, the most DE gene was AGAP028933, a small subunit ribosomal RNA which, like AAEL027000, was inconsistently expressed across replicates, and showed no Gram type-specific response in the broader data set. The other three DE genes were an odorant-binding protein (AGAP012320, *OBP25*), a thioredoxin peroxidase (AGAP011824, *TPX4*), and an aminopeptidase (AGAP013188, *APN4*). Given their functions, and their modest *padj* values, we conclude that the differential expression of these genes is likely a stochastic error. Overall, we failed to uncover conclusive evidence of a Gram type-specific transcriptional infection response in either *Ae. aegypti* or *An. gambiae* by this method. Instead, we observed that a comparison which appeared to yield robust differences in *D. melanogaster* (as a positive control for Gram type-specific transcriptional induction) yielded none in the two mosquito species.

### The intensity of the transcriptional response to bacterial infection correlates linearly with virulence and logarithmically with bacterial growth

While the genome-wide transcriptional responses to bacterial infection in *Ae. aegypti* and *An. gambiae* were not Gram type-specific, they exhibited substantial variability between different bacteria. In PCAs of the response to bacteria at 12 hours PI, the patterning was remarkably similar between the two mosquito species (Fig 5A-B). When the mean values for PC1 (accounting for 66% of variance in *Ae. aegypti*, 61% in *An. gambiae*) were plotted against each other, we obtained an R^2^ value of 0.82, indicating that the magnitude of the response to each infection (in proportion to the variability within each species across all conditions) was nearly identical between the two species (Fig 5C). We had previously observed that the order of conditions along PC1 in the complete data set (Fig 1B-C) appeared to correspond to the virulence of the infections (as measured by mosquito mortality). We had also observed that the mortality associated with all infections was closely correlated between *Ae. aegypti* and *An. gambiae* (Fig 1 S1B). Likewise, we found the mortality associated with each bacterial infection was highly correlated (R^2^ = 0.96) between the two mosquitoes (Fig 5D). We predicted that the magnitude of the transcriptional response to a given infection (as measured on PC1) would correlate with its virulence (as measured by mortality). The resulting correlations (Fig 5E-F, *Ae. aegypti* R^2^ = 0.58, *An. gambiae* R^2^ = 0.69) were modest, but censorship of *Ml* yielded larger coefficients (R^2^ = 0.77, R^2^ = 0.81, respectively). We concluded that the magnitude of the transcriptional response was related to the virulence of the pathogen for most but not all bacteria. We also hypothesized that the transcriptional variability described by PC1 was broadly reflective of the immune response. Accordingly, we plotted the total output of AMP transcripts against PC1 (Fig 5G-H) and verified close correlation (*Ae. aegypti* R^2^ = 0.90, *An. gambiae* R^2^ = 0.81) indicating that PC1 was a serviceable proxy for the relative amplitude of the immune response across conditions.

**Figure 5:**
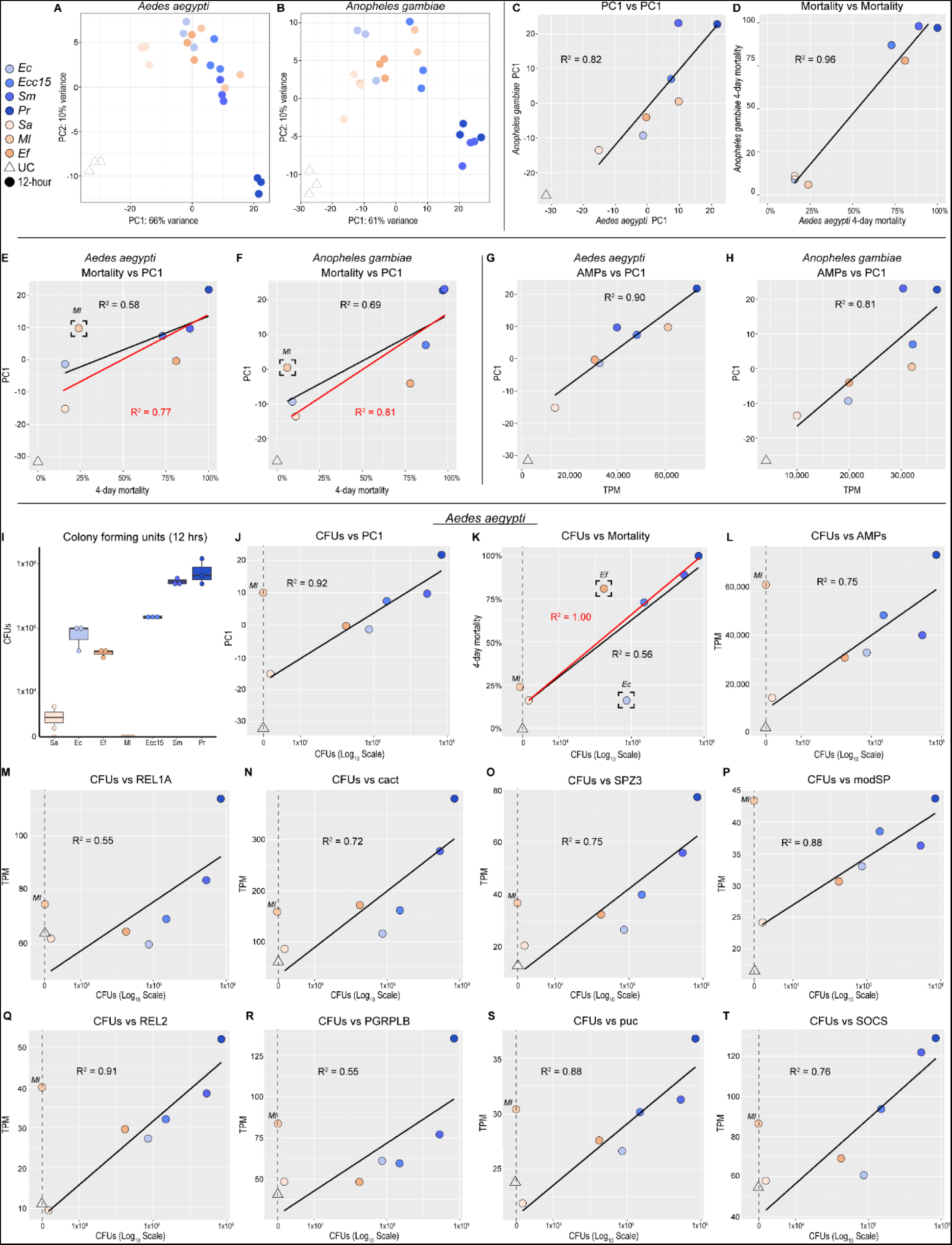
Bacterial growth correlates with hosts’ mortality, immune activation, and the amplitude of the transcriptional response to infection. Principal component analyses of transcriptomes from *Aedes aegypti* (**A**) and *Anopheles gambiae* (*s.l.*) (**B**) mosquitoes challenged with *Escherichia coli* (*Ec*), *Erwinia carotovora carotovora 15* (*Ecc15*), *Serratia marcescens* type strain (*Sm*), *Providencia rettgeri* (*Pr*), *Staphylococcus aureus* (*Sa*), *Micrococcus luteus* (*Ml*), *Enterococcus faecalis* (*Ef*) at 12 hours post-infection. Comparison of mean PC1 values in *Ae. aegypti* versus *An. gambiae* (**C**). Comparison of 4-day mortality following challenge with bacterial pathogens in *Ae. aegypti* versus *An. gambiae* (**D**). Mortality versus PC1 in *Ae aegypti* (**E**) and *An. gambiae* (**F**). Black line and R^2^ value correspond to the correlation of all data points. Red line and R^2^ value are derived from censorship of the bracketed data point. Summed expression of antimicrobial peptides (in transcripts per million, TPM) versus PC1 in *Ae. aegypti* (**G**) and *An. gambiae* (**H**). Colony forming units (CFUs) of *Sa*, *Ec*, *Ef*, *Ml*, *Ecc15*, *Sm*, and *Pr* recovered from *Ae. aegypti* 12 hours after inoculation with 3,000 CFUs (**I**). For all the following plots, data are derived from *Ae*. *aegypti* at 12 hours post-challenge and CFUs are plotted on a log_10_ scale; UC and Ml (having y values of zero) are censored from logarithmic correlations. (**J**) CFUs versus PC1. (**K**) CFUs versus mortality. Black line and R^2^ value correspond to the correlation of all non-zero data points. Red line and R^2^ value are derived from censorship of the bracketed data points. (**L**) CFUs versus summed expression of antimicrobial peptides in TPM. (**M**) CFUs versus expression of *REL1A* in TPM. (**N**) CFUs versus expression of *cact* in TPM. (**O**) CFUs versus expression of *SPZ3* in TPM. (**P**) CFUs versus expression of AAEL023229, an ortholog of *Drosophila modSP*, in TPM. (**Q**) CFUs versus expression of *REL2* in TPM. (**R**) CFUs versus expression of *PGRPLB* in TPM. (**S**) CFUs versus expression of AAEL010411, an ortholog of *Drosophila puc*, in TPM. (**T**) CFUs versus expression of *SOCS* in TPM.

As immune pathways are responsive, in part, to the presence of PAMPs, we hypothesized that the amplitude of their activation would correlate with pathogen quantity (growth) in the mosquito at the time of sacrifice. To test this hypothesis, we quantified CFUs in *Ae. aegypti* by crushing and plating at 12 hours PI (initial inoculum: 3,000 CFUs). We found that the ranking of growth rates of the different bacteria over 12 hours corresponded to their order along PC1 (Fig 5I), with the exception of *Ml* which was eliminated from the mosquito within 12 hours of injection. When CFUs were plotted against PC1 (using a logarithmic scale, Fig 5J) we found near perfect correlation (R^2^ = 0.92). It should be noted that both *Ml* and UC were necessarily excluded from this and all following correlations, as values of zero cannot be fitted to a logarithmic curve. These results support the hypothesis that immune activation is proportional to the quantity of bacteria in the mosquito, with the caveat that *Ml* is capable of stimulating a robust response (on par with *Ecc15* and *Sm*) independent of growth.

Hypothesizing that virulence would correlate with bacterial growth, we plotted CFUs versus mortality (Fig 5K). The resulting correlation was strong for some bacteria (*Sa*, *Ecc15*, *Sm*, *Pr*), but others induced mortality that was disproportionately high (*Ef*) or low (*Ec*) relative to their growth in the mosquito. We concluded that the rate of bacterial growth in the mosquito host is a contributing factor to virulence, but that other microbe-specific attributes (*e.g.*, the presence of virulence factors) may change the interaction. Alternatively, it is possible that the causal vector is reversed, and the rate of bacterial growth may be a side-effect of the virulence of infection (*i.e.*, bacteria may grow more rapidly in an ailing host).

We next evaluated the relationships between growth (CFUs) and various components, or putative components, of the immune response. The output of AMP transcripts (Fig 5L) correlated well with CFUs (R^2^ = 0.75). Expression of the terminal transcription factor of the Toll pathway (*REL1A*) was modestly correlated (R^2^ = 0.55, Fig 5M) but expression of *cactus* (a Toll modulator, Fig 5N) was better correlated (R^2^ = 0.72). We also observed some correlation between CFUs and the expression of a spaetzle cytokine (*SPZ3*), possibly involved in Toll activation (R^2^ = 0.72, Fig 5O) and a modular serine protease (*modSP*) putatively involved in the extracellular Toll signaling cascade (AAEL023229, R^2^ = 0.88, Fig 5P). We observed robust correlation (R^2^ = 0.91) between CFUs and the expression of *REL2* (Fig 5Q), the terminal transcription factor of the Imd pathway. The amidase PGRP, *PGRPLB*, which negatively regulates Imd pathway activation (63,64), showed modest correlation with CFUs (Fig 5R, R^2^ = 0.55). We saw strong correlations (R^2^ = 0.88 and 0.76) for AAEL010411 (orthologous to *D. melanogaster puc*, Fig 5S) and *SOCS* (orthologous to *D. melanogaster Socs36E*, Fig 5T) which, respectively, modulate the JNK and JAK-STAT pathways. While none of these genes have been conclusively validated as exclusive transcriptional targets of their associated pathways in *Ae. aegypti*, it is reasonable to speculate that their upregulation could be proportional to pathway activity as part of either positive (*REL1A*, *SPZ3*, *modSP*, *REL2*) or negative feedback loops (*cact*, *PGRPLB*, *puc*, *SOCS*). In such a case, the correlation of pathway components with growth (and, by extension, each other) could indicate parallel activations of these pathways in logarithmic proportion to PAMP quantity.

Overall, we interpret the results of this analysis to fit a model (Fig 5 S1) where increasing quantities of PAMPs (the product of bacterial growth) drive immune pathway activation (potentially Imd, Toll, and JAK-STAT in parallel), resulting in the upregulation of effectors. We tentatively associate virulence (mortality) with the rate of growth, with the caveat that no causality can be established from existing data. Slow-growing microbes (*e.g.*, *Sa*) are more vulnerable to elimination than their fast-growing counterparts (*e.g.*, *Pr*), with concomitant ramifications for host survival. Finally, *Ml* activates a disproportionate immune response by an undetermined mechanism, which, we hypothesize, precipitates its early elimination from the mosquito host.

### *Aedes aegypti* discriminates between bacterial and fungal pathogen-associated molecular patterns

While we found no trace of a Gram type-specific response to infection in *Ae. aegypti* and *An. gambiae*, we did note that, in PCA, transcriptomes of *Ae. aegypti* infected with the yeast *Ca* clustered separately from those of bacteria-challenged mosquitoes on PC2 (Fig 1B). To determine whether this difference in transcriptomes was indicative of a broader divide between the transcriptional response to fungi versus bacteria, we performed a second RNAseq experiment to produce transcriptional profiles for *Ae. aegypti* and *An. gambiae* injected with conidia of the filamentous fungi *Bb* and *Ma* or challenged with high doses (OD_600_ = 100) of *Ec*HK, *Ef*HK, or *Ca*HK. When we attempted to plot data from the two experiments together in a single PCA, we found, in *Ae. aegypti*, that a batch effect separated the two data sets on PC2 (Fig 6 S1A). When we instead plotted PC1 versus PC3 (Fig 6A), we observed a distinct clustering of fungal-challenged transcriptomes, separated from the cluster of bacteria-challenged transcriptomes in *Ae. aegypti*. Loadings from this PCA can be found in Table S5. No similar separation was observed in *An. gambiae* (Fig 6B). We next plotted PCAs for the heat-killed challenges alone and, in *Ae. aegypti*, observed clear separation of *Ca*HK from UC, *Ec*HK, and *Ef*HK on PC2, accounting for approximately 15% of variance between samples (Fig 6C). Loadings from this PCA can be found in Table S5. In *An. gambiae*, *Ca*HK transcriptomes appeared separated from their *Ec*HK and *Ef*HK counterparts (Fig 6D). However, the dispersion of UC transcriptomes along both PC1 and PC2 indicates that much of the transcriptional variation within the analysis could be attributed to differences between replicates rather than treatments. Overall, our PCAs provided evidence, on a genome-wide scale, for a distinct transcriptional response to fungi and to fungal PAMPs (as opposed to bacteria and bacterial PAMPs) in *Ae. aegypti*, but no clear distinction in *An. gambiae*.

**Figure 6:**
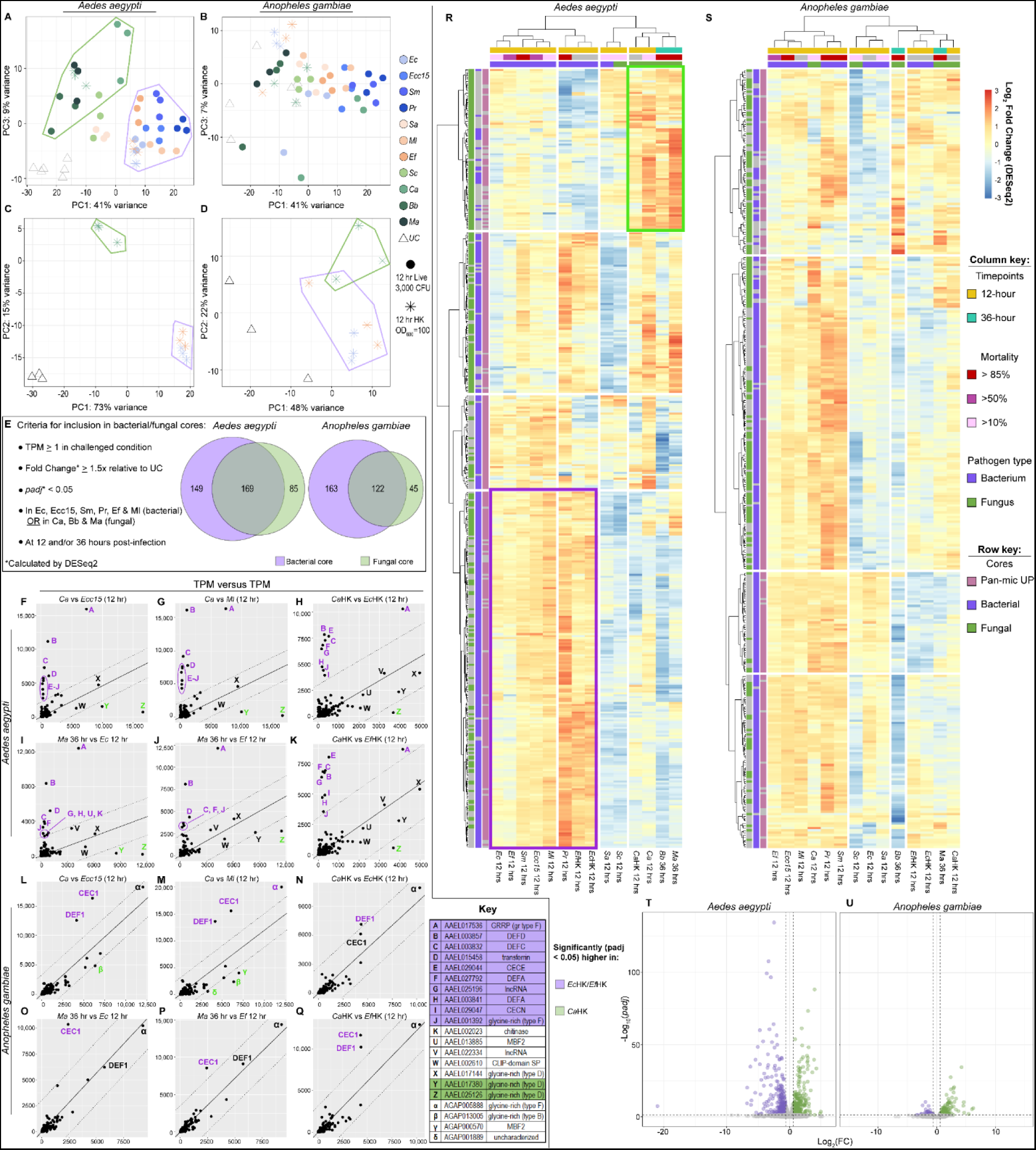
*Aedes aegypti* mounts a distinct response to fungal pathogens. Principal component analyses (PC1 vs PC3) of transcriptomes from *Ae. aegypti* (**A**) and *Anopheles gambiae* (*s.l.*) (**B**) combining data from two separate RNAseq experiments. From experiment #1: unchallenged (UC) mosquitoes and mosquitoes challenged with *Escherichia coli* (*Ec*), *Erwinia carotovora carotovora 15* (*Ecc15*), *Serratia marcescens* type strain (*Sm*), *Providencia rettgeri* (*Pr*), *Staphylococcus aureus* (*Sa*), *Micrococcus luteus* (*Ml*), *Enterococcus faecalis* (*Ef*), *Saccharomyces cerevisiae* (*Sc*), and *Candida albicans* (*Ca*). From experiment #2: unchallenged (UC) mosquitoes, and mosquitoes challenged with *Beauveria bassiana* (*Bb*), *Metarhizium anisopliae* (*Ma*), and concentrated (OD_600_ = 100) heat-killed (HK) *Ec*, *Ef*, and *Ca*; all data are from 12-hour timepoint. Principal component analyses (PC1 vs PC2) of transcriptomes from *Ae. aegypti* (**C**) and *An. gambiae* (**D**) challenged with concentrated HK *Ec*, *Ef*, and *Ca*; all data are from 12-hour timepoint. Criteria for inclusion in the bacterial and fungal cores, with Venn diagrams showing the size and relationship between cores in *Ae. aegypti* and *An. gambiae* (**E**). Plots depicting the expression (in transcripts per million) of genes in the bacterial and fungal cores in *Ae. aegypti* (**F**-**K**) and *An. gambiae* (**L**-**Q**) juxtaposing the response to *Ca* versus *Ecc15* (**F**, **L**), *Ca* versus *Ml* (**G**, **M**), *Ma* versus *Ec* (**I**, **O**), *Ma* versus *Ef* (**J**, **P**), HK *Ca* versus HK *Ec* (**H**, **N**), HK *Ca* versus HK *Ef* (**K**, **Q**); all data are from 12-hour timepoint except *Ma* which is from 36-hour timepoint. Clustering analyses of bacterial and fungal core genes in *Ae. aegypti* (**R**), and *An. gambiae* (**S**) comparing log_2_FC (calculated by DESeq2) in insects challenged with *Ec*, *Ecc15*, *Sm*, *Pr*, *Sa*, *Ef*, *Ml*, *Ca*, HK *Ca*, HK *Ec*, HK *Ef* (12-hour), and with *Bb* and *Ma* (36-hour). Volcano plots depicting genes that are differentially expressed (DESeq2 fold-change ≥ 1.5, *padj* < 0.05) in a comparison of HK *Ec*- and HK *Ef*-challenged versus HK *Ca*-challenged *Ae. aegypti* (**T**) and *An. gambiae* (**U**); all are data from 12-hour timepoint.

To search more comprehensively for a fungus-responsive signature among a narrower set of candidate genes, we defined core responses to bacterial pathogens and fungal pathogens. For each mosquito we selected all genes expressed ≥ 1 TPM that were statistically significantly (DESeq2 *padj* < 0.05) upregulated at least 1.5-fold at either 12 or 36 hours PI in *Ec*-, *Ecc15*-, *Sm*-, *Pr*-, *Ef*-, and *Ml*-challenged mosquitoes (bacterial core) or in *Ca*-, *Bb*-, and *Ma*-challenged mosquitoes (fungal core). These criteria yielded bacterial cores of 318 and 285 genes, and fungal cores of 254 and 167 genes, respectively, in *Ae. aegypti* and *An. gambiae*. In both species, the bacterial and fungal cores overlapped substantially (Fig 6E) yielding combined test sets of 403 genes (*Ae. aegypti*) and 330 genes (*An. gambiae*).

To view the expression of bacterial/fungal core genes in comparable infection conditions, we plotted their expression (in TPM) in *Ca*-infected versus *Ecc15*-infected mosquitoes (Fig 6F and L), *Ca*-infected versus *Ml*-infected mosquitoes (Fig 6G and M), *Ma*-infected versus *Ec*-infected mosquitoes (Fig 6I and O), *Ma*-infected versus *Ef*-infected mosquitoes (Fig 6J and P), *Ca*HK- versus *Ec*HK-challenged mosquitoes (Fig 6H and N), and *Ca*HK- versus *Ef*HK-challenged mosquitoes (Fig 6K and Q). We used the 12 hour PI data for all infections except for *Ma*, which elicits a stronger transcriptional response from the mosquito host at 36 hours PI (see Figs 3 S3 and S4). The pairs were selected on the basis of similarity with respect to the numbers of genes upregulated at the indicated timepoint(s) (see Fig 2A-B). After calculating the perpendicular geometric distance between each data point and a linear regression plotted through the origin, we found numerous genes at a distance greater than 2000 from the trendline. In *Ae. aegypti*, these included a set of genes that was disproportionately highly expressed in the bacteria-challenged and bacterial PAMP-challenged conditions, comprising two cecropins, three defensins, a transferrin, a long non-coding RNA, and a pair of glycine-rich type F proteins (*GRRP* and AAEL001392, see Fig 3 C-D, S2F). Below the trendline (more fungus-responsive), we observed a pair of glycine-rich type D proteins which we had previously noted in the pan-microbial core (see Fig 3 C-B, S2D). In *An. gambiae*, we found the two highly expressed AMPs from the pan-microbial core (*CEC1* and *DEF1*) were disproportionately highly expressed in the bacteria-challenged and bacterial PAMP-challenged conditions. The data points below the trendline (more fungus-responsive) in *An. gambiae* showed little consistency between comparisons. Perpendicular geometric distance scores for all genes/comparisons in this analysis can be found in Table S6. Overall, this analysis identified highly expressed genes in *Ae. aegypti* and *An. gambiae* that were disproportionately transcriptionally induced by bacterial infection (*e.g.*, cecropins, defensins, type F glycine-rich proteins) and, in *Ae aegypti* by fungal infection (type D glycine-rich proteins).

We next performed a hierarchical clustering analysis on the bacterial/fungal core genes for bacteria-, fungus-, and HK-challenged conditions (using the 36-hour timepoint for filamentous fungi, 12-hour for all others). In *Ae. aegypti* (Fig 6R) we found that the fungal conditions (live and HK) clustered separately from the bacterial conditions. We identified one set of genes that was especially responsive to fungal/fungal PAMP challenge, and a second set that was specifically responsive to bacterial/bacterial PAMP challenge. For lists of genes in these two clusters, see Table S7. In *An. gambiae* (Fig 6S) we could not identify any expression patterns specific to either bacterial or fungal challenge.

Finally, we used DESeq2 to compare *Ec*HK/*Ef*HK-challenged transcriptomes against *Ca*HK-challenged transcriptomes to determine which genes, if any, were specifically responsive to bacterial and fungal PAMPs. In *Ae. aegypti* (Fig 6T) this analysis yielded 313 genes that were higher expressed (FC ≥ 1.5, *padj* < 0.05) in *Ec*HK/*Ef*HK-challenged and 349 that were higher expressed in *Ca*HK-challenged mosquitoes. We are cautious in attributing this differential expression exclusively to a fungal versus bacterial transcriptional response. As we previously observed, patterns of differential expression in DeSeq2 comparisons between challenge conditions may be at least partially attributable to challenge-specific differences (Fig 4 S2). However, when the DE genes from the HK comparison (Table S8) were cross-referenced with PCA loadings (Fig 6A,C, Table S5), distance scores from TPM correlations (Fig 6F-K, Table S6), and clusters from the hierarchical analysis (Fig 6R, Table S7) we found many similarities, reinforcing the existence of bacteria-specific and fungus-specific transcriptional responses in *Ae. aegypti*. In *An. gambiae* (Fig 6U) we identified 61 and 175 genes that were higher expressed, respectively, in *Ec*HK/*Ef*HK-challenged and *Ca*HK-challenged mosquitoes (Table S8). Having failed to identify any bacterial or fungal-specific clusters in our hierarchical clustering analysis of the bacterial and fungal cores in *An. gambiae* (Fig 6S), we resampled the cores (Fig 6E) limiting our analysis to the genes that were differentially induced by bacterial/fungal HK challenge (Fig 6U). The resulting lists of genes were plotted in heat maps to discover whether the bacterial/fungal HK-responsive genes were, respectively, responsive to live bacterial and fungal infection. The first group (12 genes, including *DEF1*, *CEC1*, *LYSC1*, *transferrin*, and *PGRPS1*) appeared to be more robustly expressed in response to both live and HK bacterial infection versus fungal infection (Fig 6 S2A). The second group (78 genes) was less noticeably patterned by pathogen type, although we did note one CLIP-domain serine protease (*CLIPB1*) and one serpin (*SRPN4*) which appeared to exhibit modestly higher expression following fungal challenge (Fig 6 S2B). In totality, these analyses demonstrated the existence of clusters of genes specifically responsive to bacterial and fungal infection in *Ae. aegypti*, and of genes specifically responsive to bacterial infection in *An. gambiae* (see Fig 6 S3 for selected genes and expression patterns from each cohort). We were unable to conclusively demonstrate or disprove the existence of a robust fungal-specific response in *An. gambiae*.

## Material and Methods

### Mosquito Rearing

*Ae. aegypti* of the Liverpool strain were reared at Cornell University (Ithaca, New York). Larvae were reared at a density of 200 per 1 L tray from L2 stage to pupation. Each tray received 720 mg of fish food (Hikari #04428). Adults were on a diet of 10% sucrose *ad libitum*. All stages were maintained in humidified chambers at 29 °C (RH 75% ± 5%) using a 12:12-h light/dark cycle.

*An. gambiae* (*s.l.*) of the G3 strain (BEI Resources Accession # MRA-112, obtained from the Malaria Research and Reference Reagent Resource Center at CDC) were reared at Kansas State University (Manhattan, Kansas) as described previously (65). Briefly, L1 larvae were fed a slurry of 2% (w/v) baker’s yeast (Fleischmann’s Active Dry Yeast, AB Mauri, St. Louis, MO, USA). L2-L4 instar larvae were fed a slurry of 2% (w/v) ground fish food (TetraMin® Tropical Flakes, Tetra, Melle, Germany) and baker’s yeast at a 2:1 ratio. Adults were maintained on a sugar solution containing 8% fructose *ad libitum*. All stages were maintained at 27°C in a humidified chamber (RH: 75 <u>+ 5</u>%) using a 12:12-h light/dark cycle.

### Microbial culture

For *Escherichia coli, Erwinia carotovora carotovora 15, Serratia marcescens, Providencia rettgeri, Staphylococcus aureus, Micrococcus luteus, Enterococcus faecalis, Saccharomyces cerevisiae,* and *Candida albicans*: 5ml of nutrient broth (Luria Bertani media for bacteria, and Yeast Extract–Peptone–Dextrose media for yeasts) were inoculated, then incubated with shaking for about 16 hours at 200 rpm (*E. faecalis* and *M. luteus* at 37 °C and others at 29 °C). Cultures were centrifuged 20 mins at 2000 rpm, washed with 2-4 ml sterile PBS (Phosphate Buffered Saline), recentrifuged, resuspended in 1 ml sterile PBS, then diluted in sterile water to the OD_600_ corresponding to 3000 CFUs/50.6 nl (*Ec*: 0.2, *Ecc15*: 0.08, *Sm*: 0.1, *Pr*: 0.04, *Sa*: 0.06, *Ml*: 6.5, *Ef*: 0.08, *Sc*: 7.5, *Ca*: 4). Cultures were prepared at the site where injections were performed (Cornell University for *Ae. aegypti*, KSU for *An. gambiae*).

For filamentous fungi: commercial *Beauveria bassiana* (strain GHA) and *Metarhizium anisopliae* wild type (strain ARSEF Ma549) cultures and conidial suspensions were prepared as described previously (66). Briefly, *B. bassiana* and *M. anisopliae* were grown separately in Petri dishes (150 × 15 mm) containing PDA media (BD Difco, Franklin Lakes, NJ) and incubated at room temperature for 14 days. To collect conidia, sterile water and T-spreaders were used to scrape conidia from the surface of 14 petri dishes colonized by mycelia. The resulting conidial suspensions were then filtered through sterile cheesecloth into sterile conical tubes. The concentration of conidia in each suspension was determined using a hemocytometer and adjusted to a final concentration of 5.9×10^7^ conidia/ml, equivalent to 3000 conidia per injected mosquito (50.6 nl injections). The conidial suspensions were subsequently aliquoted into two microcentrifuge tubes per strain, and one tube per strain was shipped on ice from the Michel lab (KSU) to the Buchon lab (Cornell) by overnight express service. As a result, the same conidial suspensions were injected in *Ae. aegypti* and *An. gambiae* mosquitoes. Prior to injection, the viability of the conidia was assessed by placing 100 µl of the conidial suspensions (adjusted to 10^5^ conidia/ml) on PDA media. A total of 300 conidia were examined under optical microscopy (400×) 18-24 hours after inoculation, considering only conidia with germ tubes that were at least twice their diameter as viable (67). The viability of the conidia in all treatments exceeded 95% at the time of injection.

### Injections

3-day-old female mosquitoes were injected with water (mock wounding), 3000 live CFUs (bacteria and yeasts), 3000 live conidia (filamentous fungi), or HK pathogens. For HK challenges, *E. coli*, *E. faecalis* and *C. albicans* were killed by 60 min immersion of microbial suspensions at 70°C in a water bath. Heat killing was confirmed by plating. The equivalent of 3000 CFUs of each HK microbe was injected in the original experiment, and OD_600_ = 100 suspensions were injected in the follow-up experiment. A volume of 50.6 μl was injected into the thorax of each mosquito with a Nanoject injector equipped with a glass needle. Injections were performed under a stereomicroscope with carbon dioxide exposure. *Ae. aegypti* infections were performed at Cornell University; *An. gambiae* infections were performed at KSU.

### Quantification of microbial load

For CFU measurements, ten mosquitoes per condition (plus ten unchallenged mosquitoes) were collected at 12 hours after infection, surface-sterilized in 70% ethanol, dried, and individually homogenized in 1 mL sterile PBS. Samples were allowed to rest overnight at 4°C, diluted 100x, 1000x, and 10,000x in sterile PBS, then spiral plated on LB agar using a WASP II autoplate spiral plater (Microbiology International). Plates were incubated for 24 hours, or until colonies were large enough to count (*E. faecalis* and *M. luteus* at 37 °C and others at 29 °C). For microbes with distinctive colony morphology (*S. aureus*, *M. luteus*) colonies of the appropriate color/size were counted manually. For the remaining microbes, which possessed less distinguishable morphology and were seen in greater abundance, colonies were counted on plates from challenged and unchallenged mosquitoes, and the mean of the latter was subtracted from the count of each plate of the former to obtain an estimate of microbial growth. Three replicates were performed for each condition.

### RNA extraction

For RNAseq, ten mosquitoes per condition were collected and pooled at 12 hours or 36 hours after injection, per biological replicate. Three replicates were performed for each condition. All the samples homogenized in 700 μl Trizol. The samples were stored at −80 °C prior to RNA extraction via a modified phenol-chloroform method as previously described (16,42).

### Library preparation and sequencing

Libraries were prepared using the Lexogen Quantseq 3’ mRNA-seq prep kit according to the manufacturer’s instructions. Sample quality was evaluated before and after library preparation using a fragment analyzer. Libraries were pooled and sequenced on the Illumina Nextseq 500 platform using standard protocols for 75 bp single-end read sequencing at the Cornell Life Sciences Sequencing Core. Sequences have been deposited on NCBI (**accession number**).

### Data analysis pipeline

Quality control of raw reads was performed with FastQC (https://github.com/s-andrews/FastQC) (<u>Andrews et al., 2020</u>. Reads were trimmed by BBMap (https://jgi.doe.gov/data-and-tools/bbtools/) then mapped to the *Ae. aegypti* transcriptome or the *An. gambiae* transcriptome (*Aedes aegypti* LVP_AGWG *Aaeg*L5.2 and *Anopheles gambiae* PEST *Agam*P4.12, VectorBase, https://www.vectorbase.org/) (68) using Salmon version 0.9.1 with default parameters. DEseq2 (69) was used to evaluate differential expression. PCA plots and heatmaps were created using custom R scripts (available upon request). Gene ontology analysis was performed using the topGO package (classic Fisher method). The enrichment of genes categorized by Interpro domains and immune annotations was likewise evaluated using the classic Fisher method. Perpendicular geometric distances from data points to trendlines (in Figures 4 and 6) were calculated by application of Pythagorean geometry as illustrated in Fig 4 S1.

## Discussion

A primary objective of this project was to determine (a) whether mosquitoes mount a common transcriptional response to infection with a broad range of pathogens (a pan-microbial core response) and (b) to what extent such a core response is conserved between *Aedes* and *Anopheles* mosquitoes. We found that both *Ae. aegypti* and *An. gambiae* mounted a core response rich in genes with defensive functions (*e.g.,* AMPs, lysozymes, FREPs, TEPs, CLIP-domain serine proteases, pattern recognition receptors) genes related to the folding, sorting, and secretion of proteins (*e.g.*, chaperones, ER lumen protein retaining receptors, signal peptidases), and enzymes involved in macromolecule metabolism – primarily catabolism (*e.g.,* chitinases, serine proteases). The abundance of genes belonging to the various families/functions varied between the two species. For example, while the *Ae. aegypti* upregulated core included ten AMPs, the *An. gambiae* core displayed only two. Overall, however, most of the genes in each species’ upregulated core shared orthology, family, or function with one or more genes in the other species’ core, indicating that the core response to infection has remained highly conserved over 160 million years (70) of evolution.

Among the genes in the pan-microbial cores of *Ae. aegypti* and *An. gambiae*, we observed several genes/gene categories which were upregulated upon infection, but whose function in the context of infection is unknown or poorly characterized. One such category was the MBF2 factors, four of which were upregulated in the *Ae. aegypti* core, one in the *An. gambiae* core. In mosquitoes, the precise biological function of these factors has yet to be elucidated. However, we have previously observed that, under basal conditions, several of these factors are strongly and specifically expressed in the mosquito proventriculus and anterior midgut, which are also highly enriched for AMP expression (42). We hypothesize that MBF2 factors may play a role in promoting the expression of immune effectors in the conditions and compartments where they are called for.

The upregulated cores of both species also contained a plurality (three in *Ae. aegypti*, two in *An. gambiae*) of farnesoic O-methyl transferases, an enzyme which was shown to catalyze the rate-limiting step in the synthesis of juvenile hormone III in the cockroach *Diploptera punctata* (47). This is intriguing, as juvenile hormone is not a canonical part of insects’ response to infection and has, indeed, been reported to suppress infection resistance (71–74). Subsequent research has failed to demonstrate any role for the enzyme in the synthesis of juvenile hormone in *D. melanogaster* (75). It would be interesting to repeat this assessment in *Ae. aegypti* or *An. gambiae*, or to silence these enzymes to determine what role, if any, they play in the outcome of infection. It might also be interesting to undertake a systematic examination of how widespread the transcriptional upregulation of these enzymes is among arthropods during infection, as they have previously been noted in the transcriptional infection responses of *Bombyx mori* (76) and of *Exopalaemon carinicauda*, the white-tailed prawn (77).

Uncharacterized glycine-rich proteins were prominent in the core infection responses of both species. We further noted that the level of glycine overrepresentation among some of these genes rivaled the glycine content of known AMPs and, further, that some exhibited other characteristics of AMPs (high isoelectric points, signal peptide sequences, etc.). A comparison of the sequences of glycine-rich proteins in both cores uncovered six groups (of two or more genes), which were shared in both species’ upregulated cores. Among these, two types (D and F) were of special interest. The type D glycine-rich proteins comprised one gene in *An. gambiae* and six in *Ae. aegypti* with highly conserved sequences. In a review of the literature, we discovered that several members of this family have been found to be transcriptionally responsive to infection in other contexts. The *An. gambiae* gene, AGAP001508, responds to *Plasmodium* infection (78), and several of the *Ae. aegypti* genes (AAEL025126, AAEL025531, and AAEL017380) have variously been found to be upregulated by chikungunya and dengue fever virus infection (79–81). Our own results showed that four of the six *Ae. aegypti* genes were robustly, and specifically, upregulated by fungal infection (see Fig 6 S3). The type F glycine-rich proteins included the AMP holotricin (*GRRP*) which is the highest-expressed bacterial core gene for most of the 12-hour bacterial challenge conditions we examined (see Fig 4 H, K, M). The *An. gambiae* type F glycine-rich protein, AGAP005888, was the highest expressed bacterial core gene in the same conditions (see Fig 4 I, L, N). In a review of the literature, we also found that the unnamed *Ae. aegypti* type F glycine-rich protein, AAEL001392, was significantly upregulated by dengue fever virus infection (81,82). In the light of our own transcriptomic data, the transcriptomic data of other researchers, and the physiochemical properties of the proteins in question, we strongly suspect that type D and type F glycine-rich proteins are antimicrobial effectors of some kind, and we propose to explore their efficacy against different types of mosquito pathogens at some later date.

This study’s main objective was to contrast the transcriptomes of mosquitoes following infection with Gram-negative, Gram-positive, and fungal pathogens to inform a model for the mechanisms governing the transcriptional response to each. The canonical model for pathogen type-specific immune regulation in insects, as described in *D. melanogaster*, calls for the activation of the Imd pathway in response to Gram-negative PAMPs, and the activation of the Toll pathway in response to Gram-positive and fungal PAMPs (Fig 6 S4B). We found this model inconsistent with the results of our study. An exhaustive comparison of the transcriptomes of mosquitoes challenged with Gram-negative and Gram-positive pathogens (live and HK) failed to uncover any evidence of distinct transcriptional programs responsive to Gram type. Rather, we found that the variance in the transcriptional response to bacterial infection was not in kind, but in amplitude, which was positively correlated with virulence and the rate of bacterial growth (with the noted exception of *M. luteus*). By contrast, comparisons of the transcriptomes of bacteria-challenged versus fungus-challenged mosquitoes (live and HK) yielded clear evidence of separate transcriptional programs in both *Ae. aegypti* and *An. gambiae*. In both species, sharp upregulations of cecropins and defensins were observed following bacterial infection, with fungal infections of similar or greater virulence mustering far less expression (Fig 6 S3). In *Ae. aegypti*, we were also able to observe a consistent fungus-specific response including (among other things) a pair of highly expressed glycine-rich proteins (type D) (Fig 6 F, G, I). While we were unable to identify a similarly robust fungus-specific response in *An. gambiae*, possibly due to greater variability between replicates and lower statistical power, the presence of a bacterial-specific response in this species is sufficient to demonstrate discrimination between bacteria and fungi in this species’ transcriptional infection response.

With the fact of separate transcriptional programs in mosquitoes for bacterial versus fungal infection established, we can only speculate as to the precise mechanisms underlying them. As compared to *D. melanogaster*, the most parsimonious model in terms of change would be a peptidoglycan-responsive Imd pathway (Gram type-independent) and a β-glucan-responsive Toll pathway (Fig 6 S4C). In a variation of this scenario, Imd might be responsive to peptidoglycan while Toll is activated slightly by peptidoglycan and/or host damage and more robustly by fungal PAMPs. Alternatively, both Imd and Toll might contribute in parallel or, as has been proposed previously (83), synergistically to the activation of AMP expression in response to bacterial infection, while some undetermined other pathway senses and responds specifically to fungal infection (Fig 6 S4D). In a third scenario, Imd and Toll are activated simultaneously by both bacteria and fungi, responding more robustly to the former (mediating the expression of cecropins and defensins) while an unknown fungus-responsive pathway mediates the upregulation of the small cohort of fungus-responsive genes (type D glycine-rich proteins) (Fig 6 S4E). These scenarios are not exhaustive. We could also envision models of greater complexity (*e.g.*, universal activation of the Toll pathway through damage-sensing, coupled with Imd-mediated repression of Toll targets) which could also explain the transcriptional patterns uncovered in this study. More work is required to determine which scenario best describes the relationship between pathogen, pathway, and transcriptional targets in mosquitoes. An examination of the expression patterns of putative Toll and Imd targets and pathway-related genes in our data set yielded no concrete conclusion on this point (Fig 6 S4A). We will note that one persuasive piece of evidence in favor of the Toll dependence of the fungal-specific response here described is the highly significant upregulation of the type-D glycine-rich proteins AAEL017380, AAEL025531, AAEL025126, and AAEL021929 in the whole bodies of female *Ae. aegypti* following RNAi-mediated knockdown of the repressor *cactus* documented by Sneed et. al (84).

## Conclusion

In this manuscript we analyzed the transcriptional responses of *Ae. aegypti* and *An. gambiae* mosquitoes to systemic infection with Gram-negative bacteria, Gram-positive bacteria, yeasts, and filamentous fungi, and to systemic challenge with heat-killed bacterial and fungal pathogens. We demonstrate that each mosquito mounts a core transcriptional response to all types of infection, and that this response is well conserved between the *anopheline* and *culicine* lineages with respect to both function and orthology. An exhaustive search for Gram type-specific responses in both species yielded no results, but comparisons of bacteria-challenged versus fungus-challenged transcriptomes revealed that both *Ae. aegypti* and *An. gambiae* discriminate in their transcriptional responses to bacteria and fungi. This work demonstrates that mosquitoes’ systemic immune responses do not follow the canonical model described in *D. melanogaster*, and sets the stage for future investigations of immune regulation in these crucial vector species.

## Supporting information

Supplemental Table 1 pan-microbial core genes (values)

Supplemental Table 2 updated immune gene list (values)

Supplemental Table 3 GO interpro & defense gene enrichment

Supplemental Table 4 distance scores TPM correlations G− vs G+

Supplemental Table 5 PCA loadings

Supplemental Table 6 distance scores TPM correlations bacteria vs fungi

Supplemental Table 7 Aedes bacteria & fungi clustered genes

Supplemental Table 8 HK DESeq genes

## Acknowledgements

We thank Dr. Patil Tawidian (Kansas State University) for the production of *Metarhizium anisopliae* and *Beauveria bassiana* spores and all members of the Michel lab for *An. gambiae* rearing. We would also like to thank Dr. Erika Mudrak for invaluable assistance with custom scripts and statistical analyses.

## Supplemental Figures and Legends

**Fig 1 S1:**
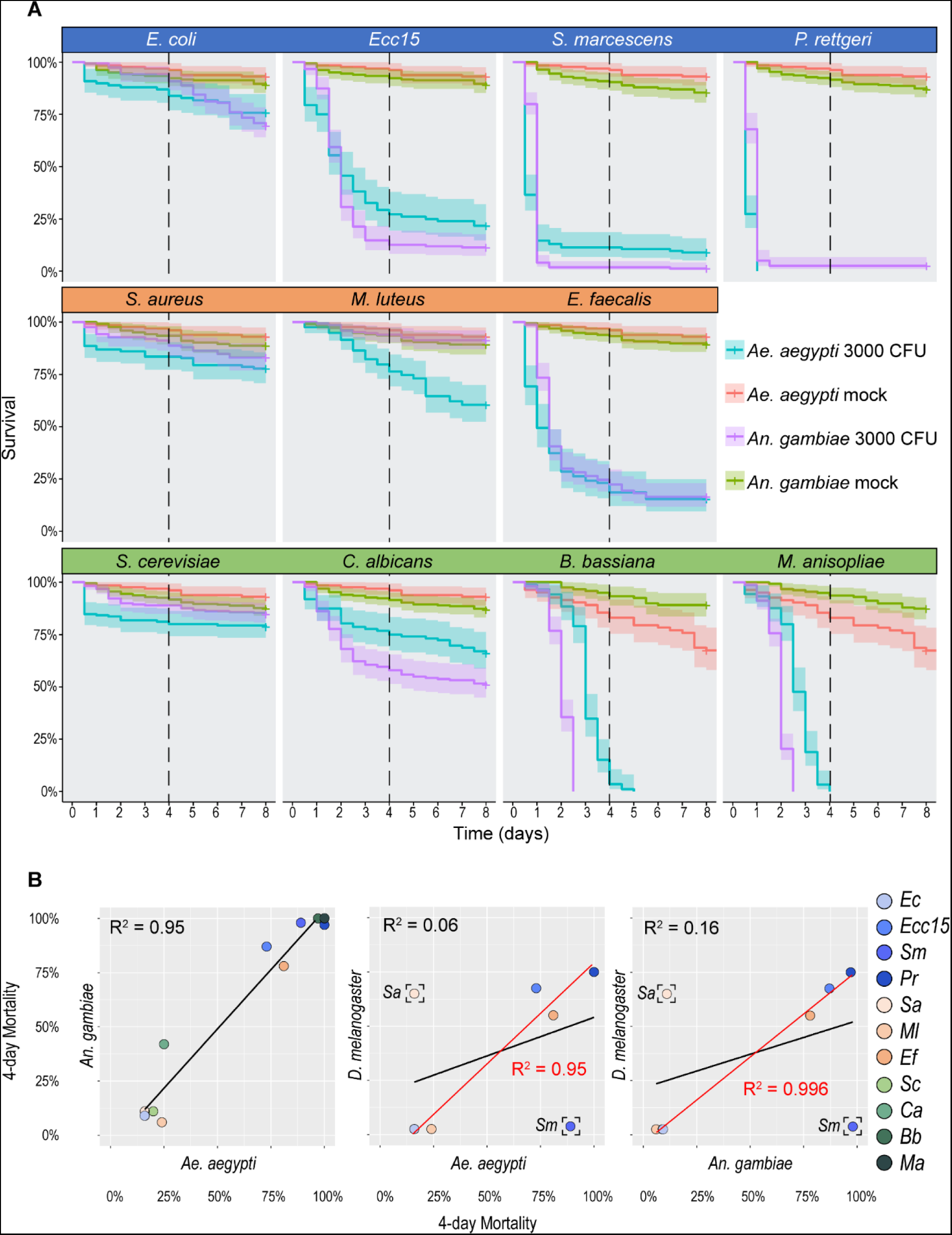
Mortality following systemic infection with bacteria, yeasts, and filamentous fungi is closely correlated in *Aedes aegypti* and *Anopheles gambiae*. (**A**) Mosquitoes were injected with 3,000 colony-forming units or conidia of Gram-negative bacteria (blue row), Gram-positive bacteria (orange row), and fungal (green row) pathogens. Pathogens include *Escherichia coli* (*Ec*), *Erwinia carotovora carotovora 15* (*Ecc15*), *Serratia marcescens* type strain (*Sm*), *Providencia rettgeri* (*Pr*), *Staphylococcus aureus* (*Sa*), *Micrococcus luteus* (*Ml*), *Enterococcus faecalis* (*Ef*), *Saccharomyces cerevisiae* (*Sc*), *Candida albicans* (*Ca*), *Beauveria bassiana* (*Bb*), and *Metarhizium anisopliae* (*Ma*). Dotted lines mark 4 days post-infection; in this study, mortality at 4 days serves as a proxy value for pathogen virulence in a host. (**B**) Correlations of mortality of all pathogens in *Ae. aegypti* versus *An. gambiae* (*s.l.*), *Ae. aegypti* versus *Drosophila melanogaster*, and *An. gambiae* versus *D. melanogaster*. Bracketed data points are included in the linear regressions shown in black, but censored from the linear regressions shown in red.

**Fig 2 S1:**
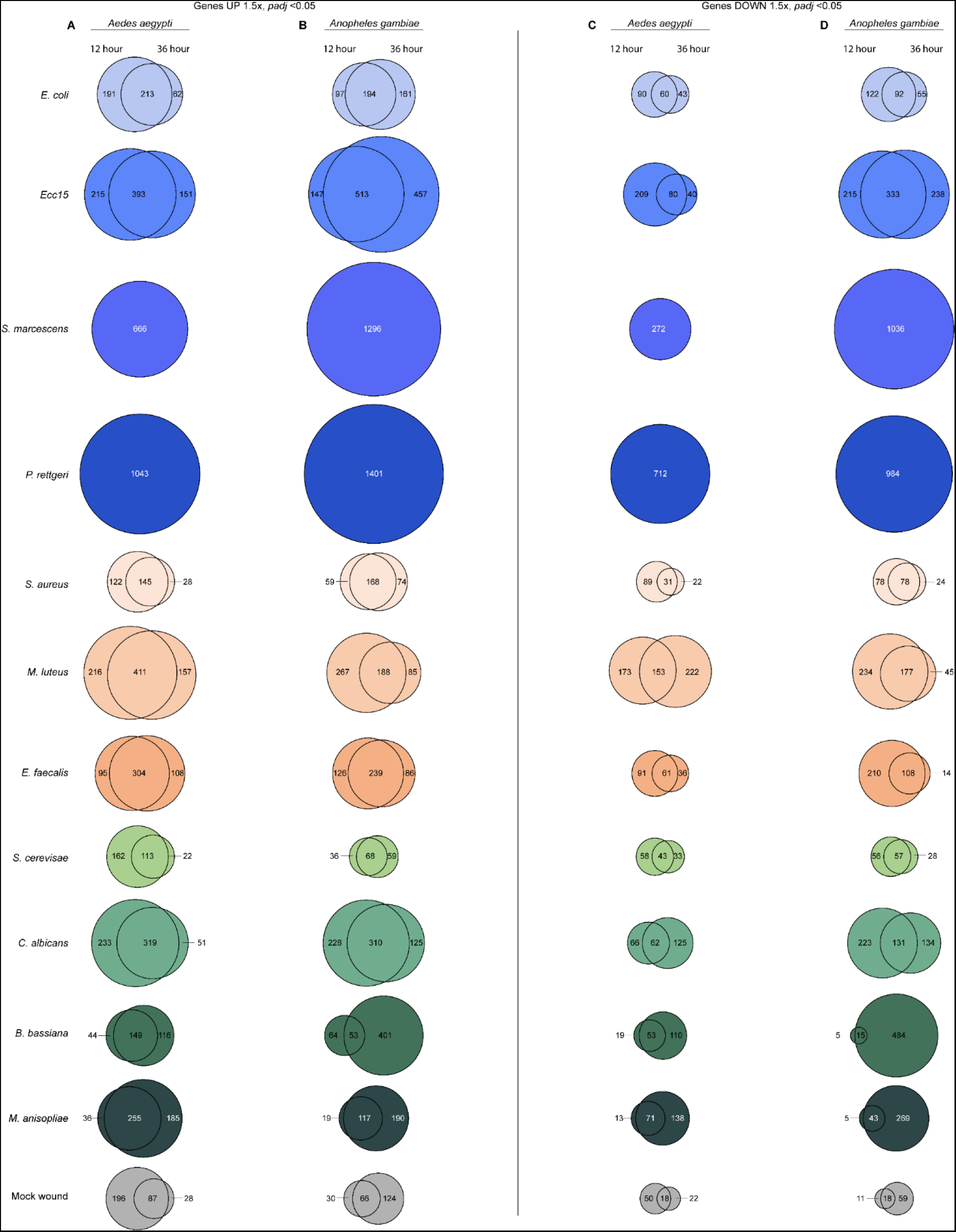
Differentially expressed genes in *Aedes aegypti* and *Anopheles gambiae* at 12 versus 36 hours post-infection. Venn diagrams displaying the number of genes upregulated (left) and downregulated (right) at 12 hours versus 36 hours post-challenge in *Ae. aegypti* (**A**, **C**) and *An. gambiae* (*s.l.*) (**B**, **D**) mosquitoes. Challenges included mock wounding and live infection with 3,000 CFUs of *Escherichia coli*, *Erwinia carotovora carotovora 15* (*Ecc15*), *Serratia marcescens* type strain, *Providencia rettgeri*, *Staphylococcus aureus*, *Micrococcus luteus*, *Enterococcus faecalis*, *Saccharomyces cerevisiae*, *Candida albicans*, and 3000 conidia of *Beauveria bassiana,* and *Metarhizium anisopliae*. Transcriptomes were assayed by RNAseq at 12 and 36 hours post-challenge, except where mortality was too high to collect the later timepoint (*S. marcescens* and *P. rettgeri*). Criteria for differential expression are ≥ 1.5x fold-change (up) or −1.5x fold-change (down) relative to unchallenged, and *padj* < 0.05, as calculated by DESeq2.

**Fig 2 S2:**
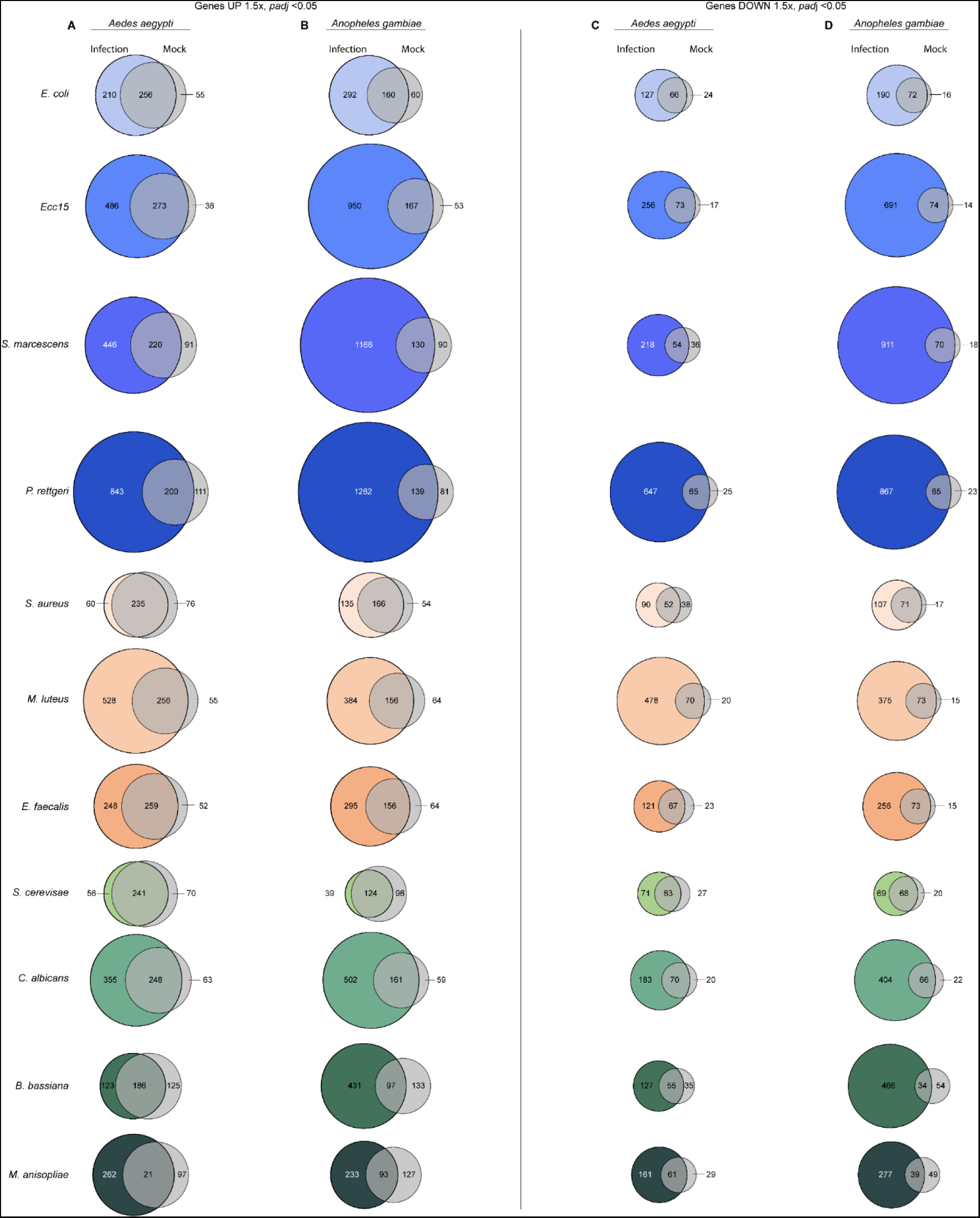
Differentially expressed genes in *Aedes aegypti* and *Anopheles gambiae*, infection versus mock wounding. Venn diagrams display the number of genes upregulated (left) and downregulated (right) in infected *Ae. aegypti* (**A**, **C**) and *An. gambiae* (*s.l.*) (**B**, **D**) mosquitoes versus mock-wounded counterparts. Live challenges included infection with 3,000 CFUs of *Escherichia coli*, *Erwinia carotovora carotovora 15* (*Ecc15*), *Serratia marcescens* type strain, *Providencia rettgeri*, *Staphylococcus aureus*, *Micrococcus luteus*, *Enterococcus faecalis*, *Saccharomyces cerevisiae*, *Candida albicans*, and 3000 conidia of *Beauveria bassiana,* and *Metarhizium anisopliae*. Transcriptomes were assayed by RNAseq at 12 and 36 hours post-challenge, except where mortality was too high to collect the later timepoint (*S. marcescens* and *P. rettgeri*). The count of regulated genes per condition is inclusive of both timepoints. Criteria for differential expression are ≥ 1.5x fold-change (up) or −1.5x fold-change (down) relative to unchallenged, and *padj* < 0.05, as calculated by DESeq2.

**Fig 2 S3:**
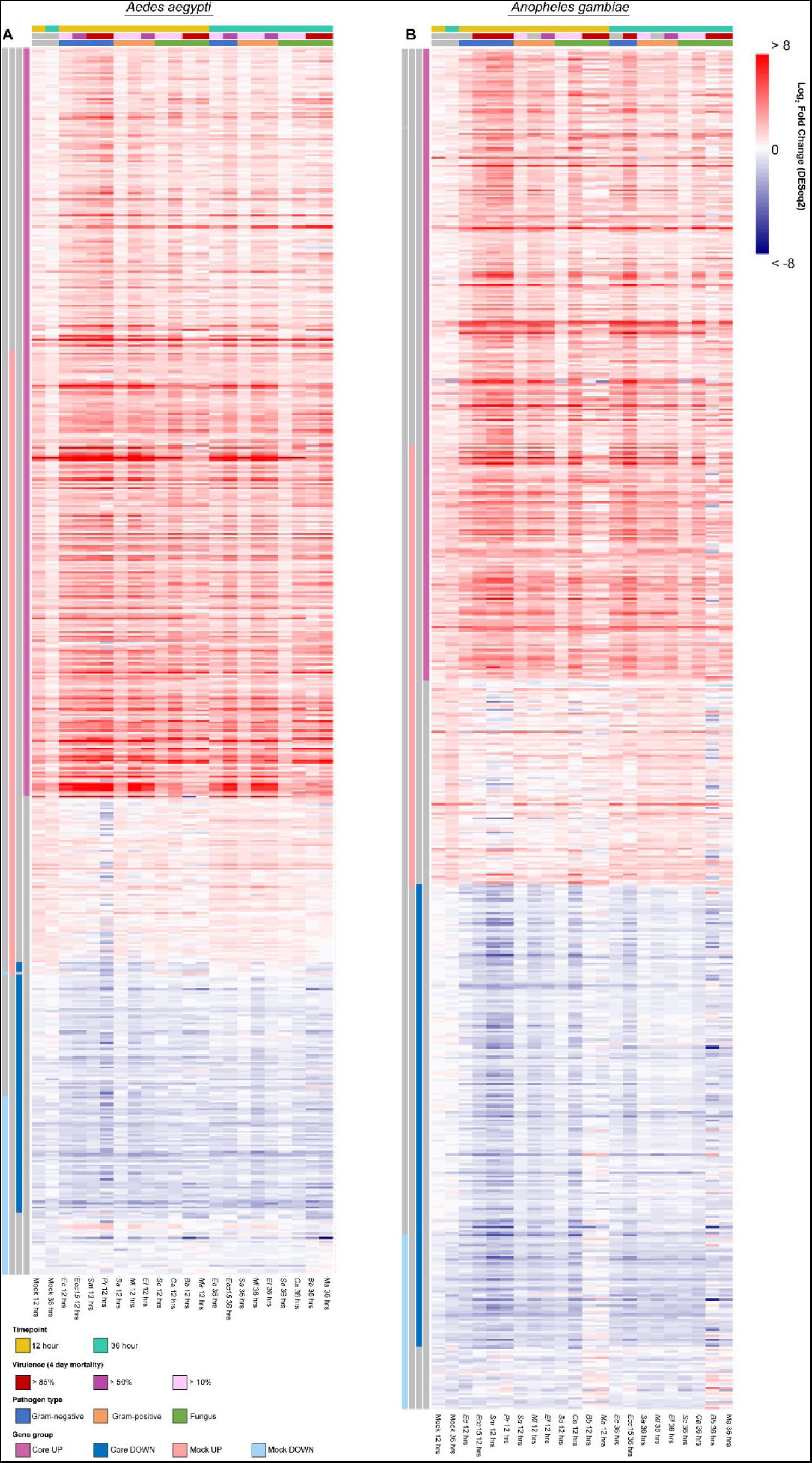
The amplitude of change in pan-microbial core genes is greater following live infection compared with mock wounding. Unscaled heatmaps of fold change (relative to unchallenged, calculated by DESeq2) in mock-infected and live-infected conditions at 12 and 36 hour post-challenge in *Aedes aegypti* (**A**) and *Anopheles gambiae* (*s.l.*) (**B**) mosquitoes, including all genes from the following groups: UP core genes, DOWN core genes, and genes that were differentially expressed (at either timepoint) following mock infection. Criteria for differential expression are ≥ 1.5x fold-change (up) or −1.5x fold-change (down) relative to unchallenged, and *padj* < 0.05, as calculated by DESeq2.

**Fig 3 S1:**
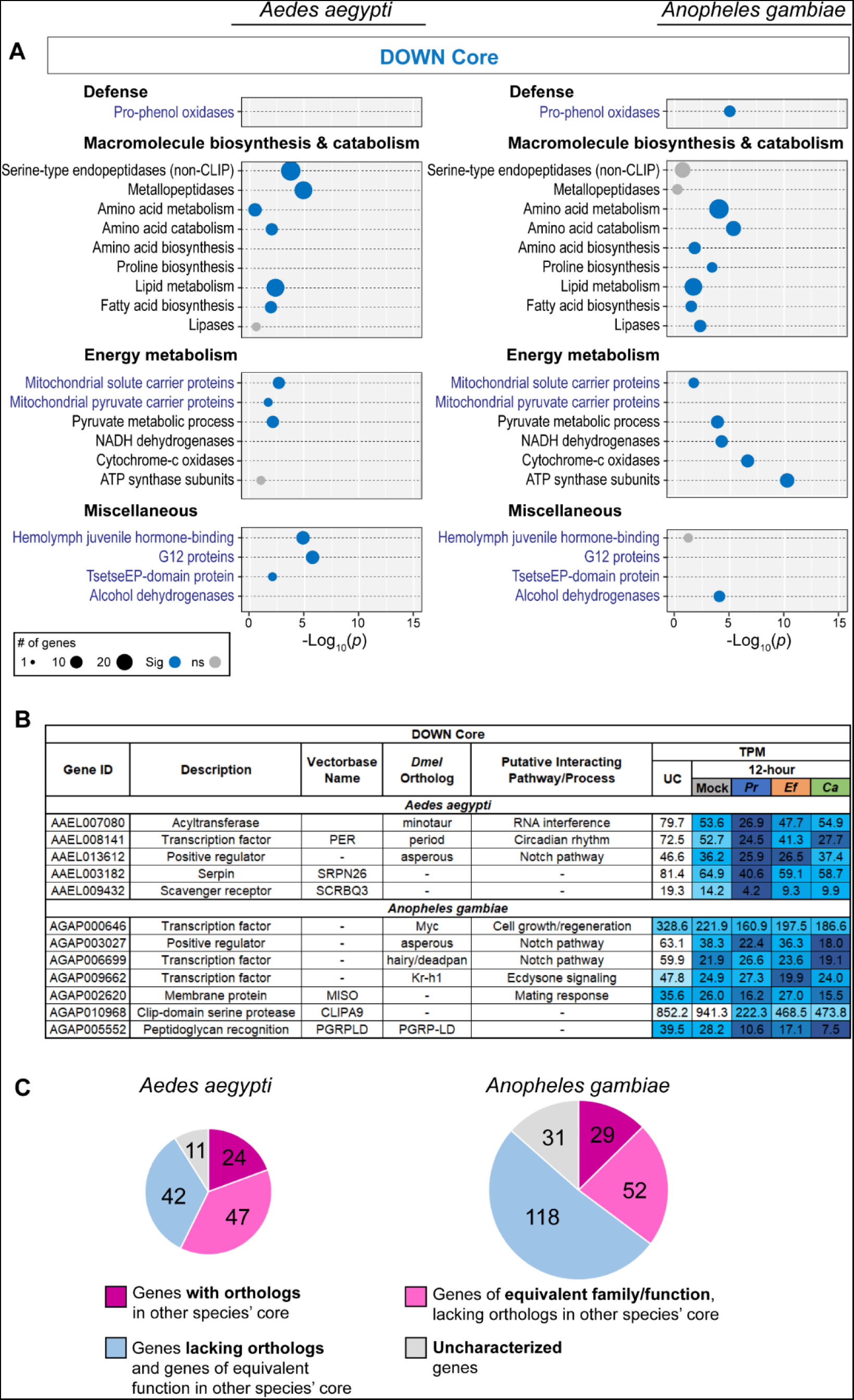
The downregulated core response to infection in *Aedes aegypti* and *Anopheles gambiae*. Bubble plots (**A**) display categories of genes enriched in the downregulated pan-microbial cores of *Ae. aegypti* and/or *An. gambiae*. Black text indicates category is from TopGO. Blue text indicates custom category (either from immune gene list, or defined by the presence of InterPro domain(s)). The size of the bubble is proportional to the number of genes from the given category in the downregulated core. The placement of the bubble along the x axis corresponds to the statistical significance of the enrichment (Fisher’s exact test). Gray bubbles indicate *p* value ≥ 0.05 (not significant enrichment). (**B**) Table of selected genes in the downregulated cores of *Ae. aegypti* and *An. gambiae*. (**C**) Pie charts describing the orthology and functional similarities shared by the *Ae. aegypti* and *An. gambiae* downregulated cores.

**Fig 3 S2:**
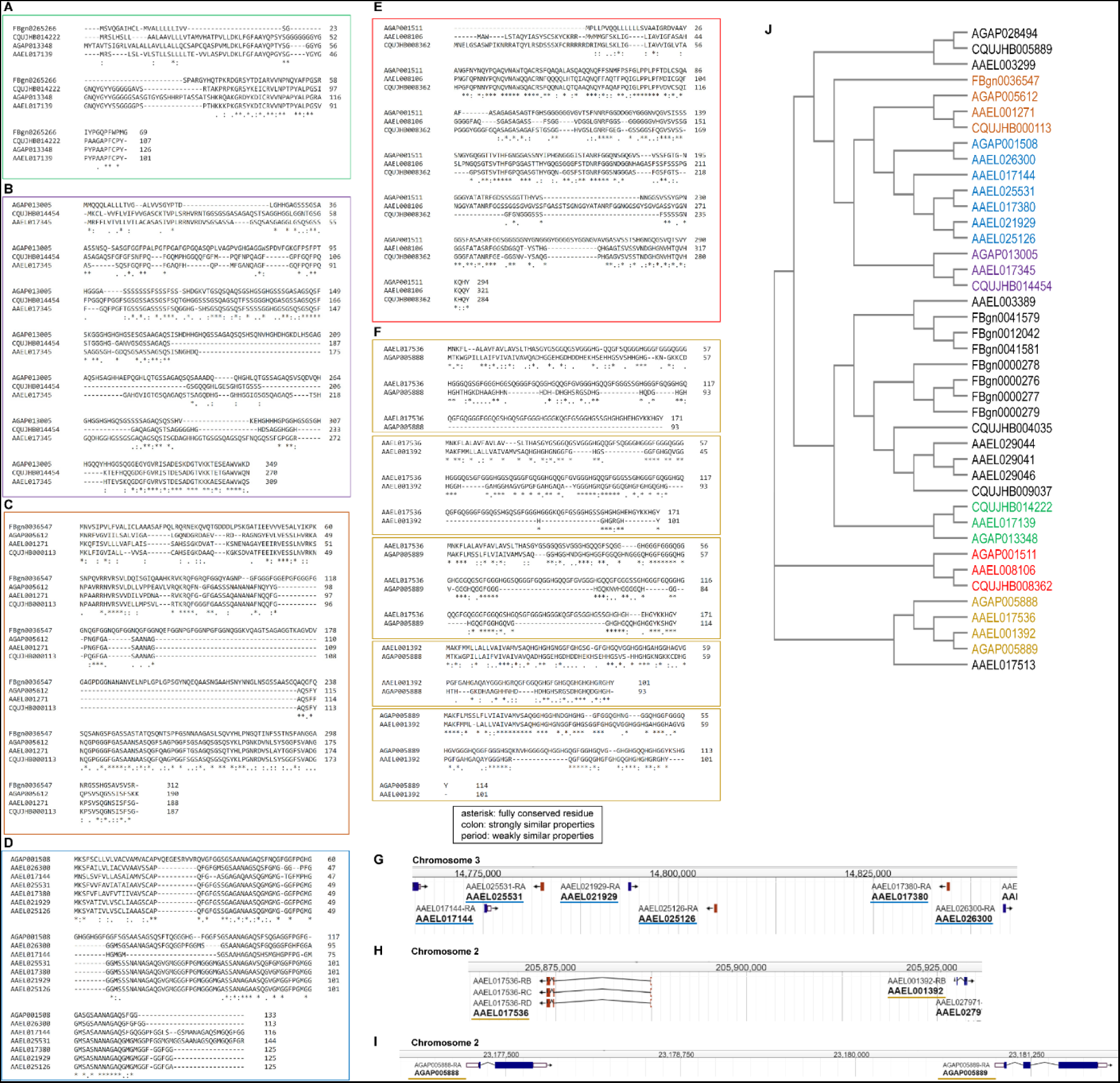
Infection-responsive glycine-rich proteins in *Aedes aegypti* and *Anopheles gambiae* share sequence similarity with each other, and with genes from *Culex quinquefasciatus* and *Drosophila melanogaster*. (**A**-**F**) Sequence alignments of glycine-rich (>13%) proteins by Clustal Omega. (**G-I**) Genomic locations of glycine-rich proteins of clusters D and F in *Ae. aegypti* and *An. gambiae*. Images are derived from VectorBase Genome Browser. (**J**) A dendrogram of glycine-rich proteins from the *Ae. aegypti* and *An. gambiae* cores, together with related proteins from *Drosophila melanogaster* and *Culex quinquefasciatus*, generated by Clustal Omega.

**Fig 3 S3:**
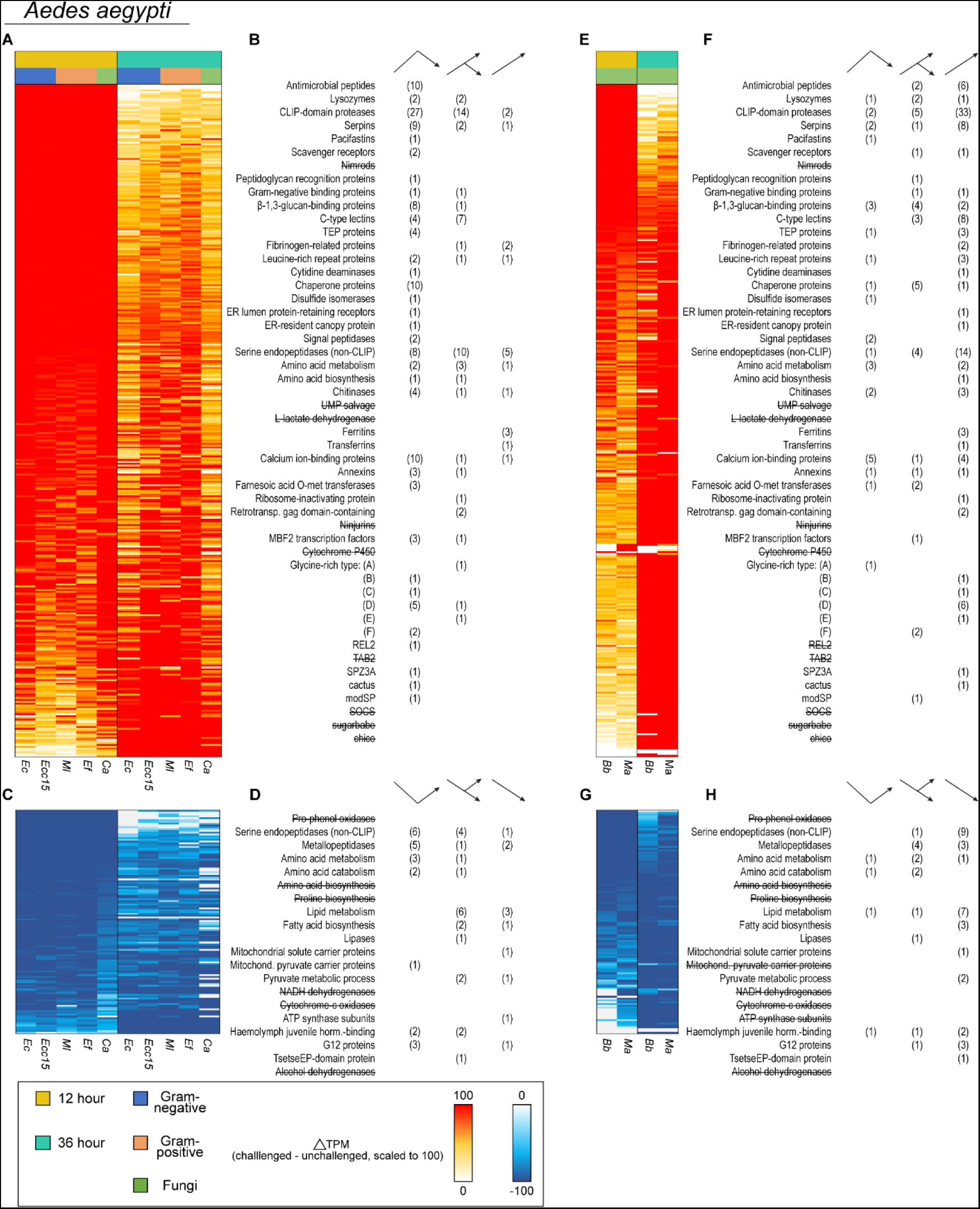
The core transcriptional response is implemented more rapidly following systemic infection with bacteria and yeast versus with filamentous fungi in *Aedes aegypti*. Heatmaps of change in expression (TPM difference in challenged versus unchallenged) of upregulated (**A**, **E**) and downregulated (**C**, **G**) pan-microbial core genes in *Ae. aegypti* challenged with *Escherichia coli* (*Ec*), *Erwinia carotovora carotovora 15* (*Ecc15*), *Micrococcus luteus* (*Ml*), *Enterococcus faecalis* (*Ef*), and *Candida albicans* (*Ca*) (**A**, **C**) or with *Beauveria bassiana* (*Bb*) and *Metarhizium anisopliae* (*Ma*) (**E**, **G**). Values are scaled by infection and grouped by timepoint. Tallies of genes in the descending (peaked before 36 hours for at least 4/5 of conditions for bacterial/yeast infections or for 2/2 of conditions for filamentous fungi infections) ascending (peaked after 36 hours for at least 4/5 conditions for bacterial/yeast infections or for 2/2 of conditions for filamentous fungal infections), and intermediate (met neither of the previous criteria) cohorts of the upregulated (**B**, **F**) and downregulated (**D**, **H**) core genes in mosquitoes challenged with *Ec*, *Ecc15*, *Ml*, *Ef*, and *Ca* (**B**, **D**) or with *Bb* and *Ma* (**F**, **H**). Genes that were not regulated at least 1.5-fold in both *Bb* and *Ma* were excluded from the counts in **F** and **H**. Categories with a null count for the given species/infection type are shown with a strike-through font.

**Fig 3 S4:**
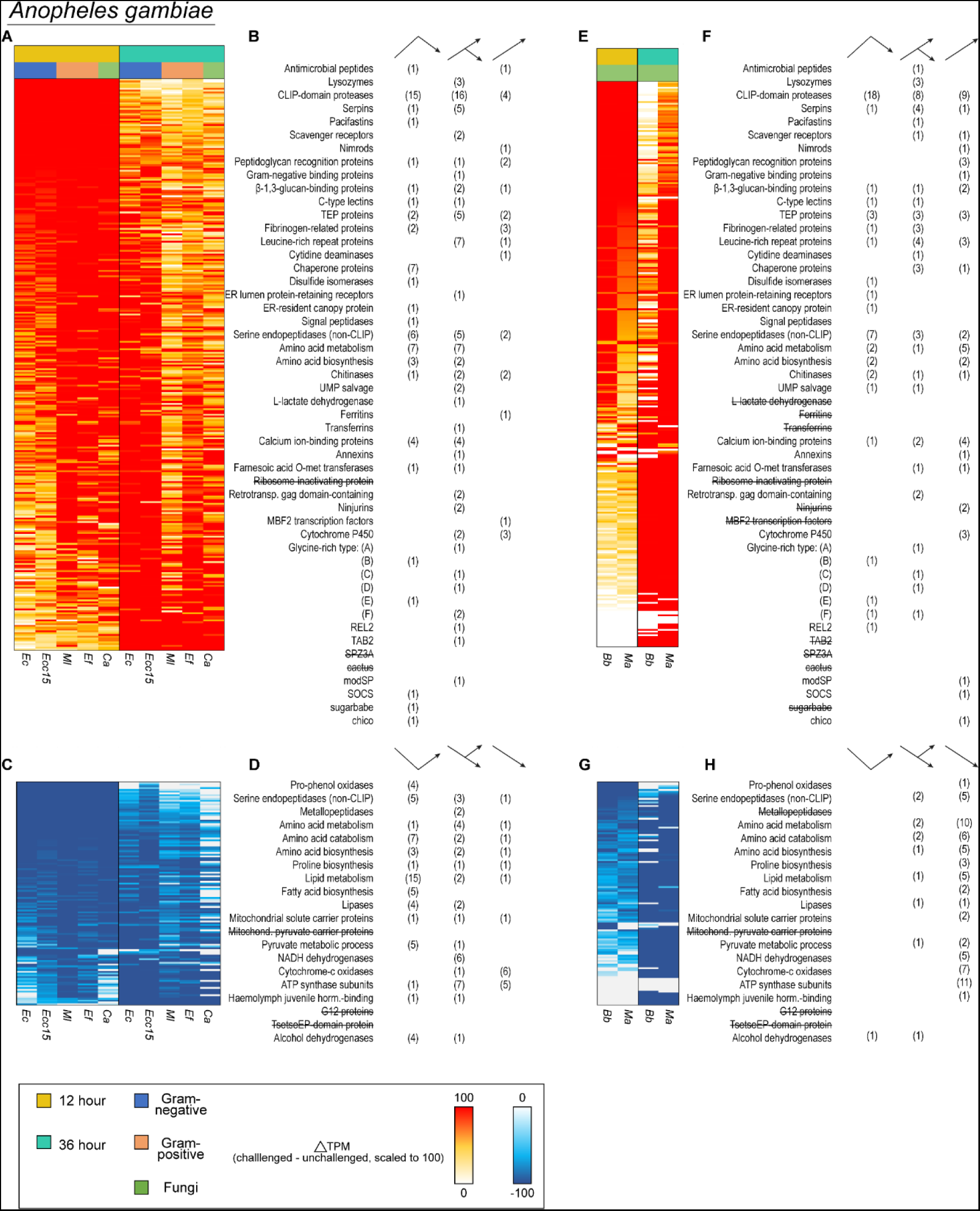
The core transcriptional response is implemented more rapidly following systemic infection with bacteria and yeast versus with filamentous fungi in *Anopheles gambiae*. Heatmaps of change in expression (TPM difference in challenged versus unchallenged) of upregulated (**A**, **E**) and downregulated (**C**, **G**) pan-microbial core genes in *An. gambiae* (*s.l*.) challenged with *Escherichia coli* (*Ec*), *Erwinia carotovora carotovora 15* (*Ecc15*), *Micrococcus luteus* (*Ml*), *Enterococcus faecalis* (*Ef*), and *Candida albicans* (*Ca*) (**A**, **C**) or with *Beauveria bassiana* (*Bb*) and *Metarhizium anisopliae* (*Ma*) (**E**, **G**). Values are scaled by infection and grouped by timepoint. Tallies of genes in the descending (peaked before 36 hours for at least 4/5 of conditions for bacterial/yeast infections or for 2/2 of conditions for filamentous fungi infections) ascending (peaked after 36 hours for at least 4/5 conditions for bacterial/yeast infections or for 2/2 of conditions for filamentous fungi infections), and intermediate (met neither of the previous criteria) cohorts of the upregulated (**B**, **F**) and downregulated (**D**, **H**) core genes in mosquitoes challenged with *Ec*, *Ecc15*, *Ml*, *Ef*, and *Ca* (**B**, **D**) or with *Bb* and *Ma* (**F**, **H**). Genes that were not regulated at least 1.5-fold in both *Bb* and *Ma* were excluded from the counts in **F** and **H**. Categories with a null count for the given species/infection type are shown with a strike-through font.

**Fig 4 S1:**
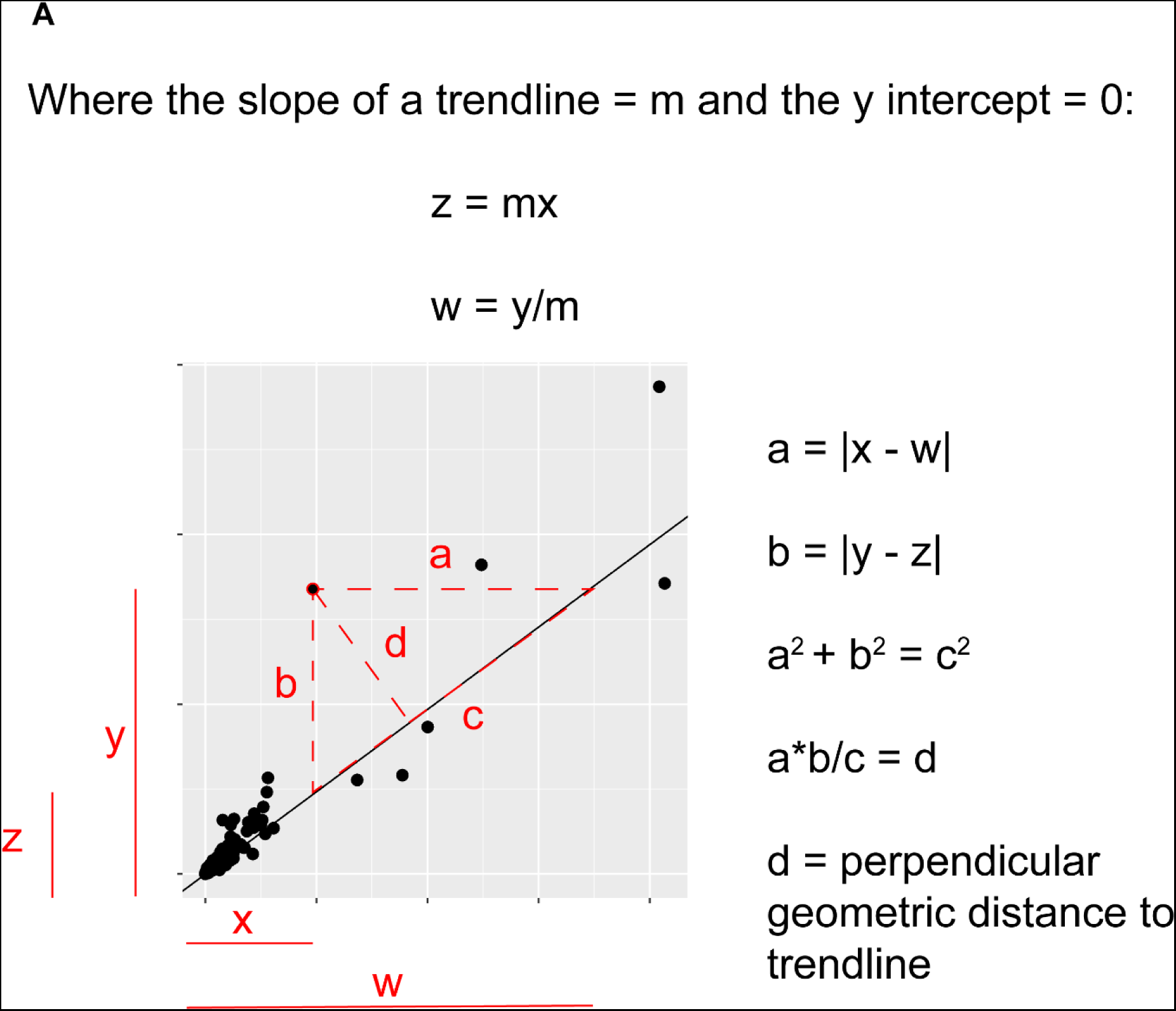
Calculation of the geometric perpendicular distance of a data point from a trendline. Method for deriving geometric distances using x and y values and the slope of a linear regression.

**Figure 4 S2:**
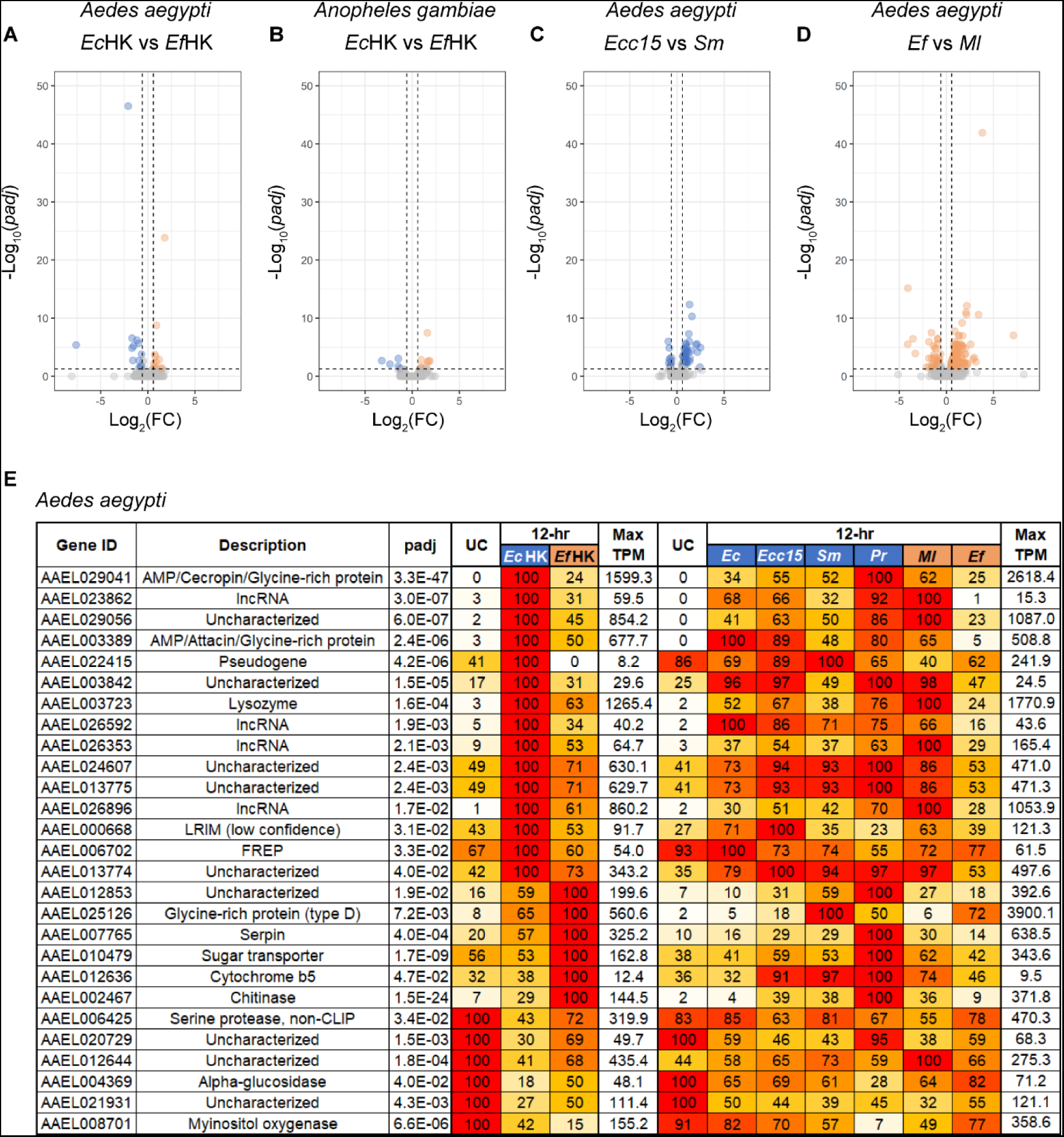
One-to-one comparisons of bacteria-challenged transcriptomes fail to demonstrate Gram type-specific transcriptional responses. Volcano plots depicting genes that are differentially expressed (DESeq2 fold-change ≥ 1.5, *padj* < 0.05) in a comparison of heat-killed *Escherichia coli* (*Ec*HK) versus heat-killed *Enterococcus faecalis* (*Ec*HK) (OD_600_ = 100) in *Aedes aegypti* (**A**) and *Anopheles gambiae* (**B**); further volcano plots compare the transcriptomes of *Ae. aegypti* challenged with live (3000 CFU) *Erwinia carotovora carotovora 15* (*Ecc15*) versus *Serratia marcescens* type strain (*Sm*) (**C**) and *Enterococcus faecalis* versus (*Ef*) *Micrococcus luteus* (*Ml*) (**D**). (**E**) Table comparing TPMs of differentially expressed genes from panel (**A**) in live and heat-killed challenged conditions in *Ae. aegypti*. Conditions on the left-hand side include unchallenged mosquitoes (UC) and mosquitoes challenged with heat-killed *Ec* (*Ec*HK), heat-killed *Ef* (*Ef*HK) (OD_600_ = 100); all data are from the second RNAseq experiment. The right-hand side compares unchallenged (UC), and challenged with live *Escherichia coli* (*Ec*), *Erwinia carotovora carotovora 15* (*Ecc15*), *Serratia marcescens* type strain (*Sm*), *Providencia rettgeri* (*Pr*), *Micrococcus luteus* (*Ml*), and *Enterococcus faecalis* (*Ef*). Within each comparison data are scaled to show relative expression so that the condition with the highest expression (shown in the max TPM column) is scored at ‘100’ and all lower expression values are expressed as a percentage of 100.

**Fig 5 S1:**
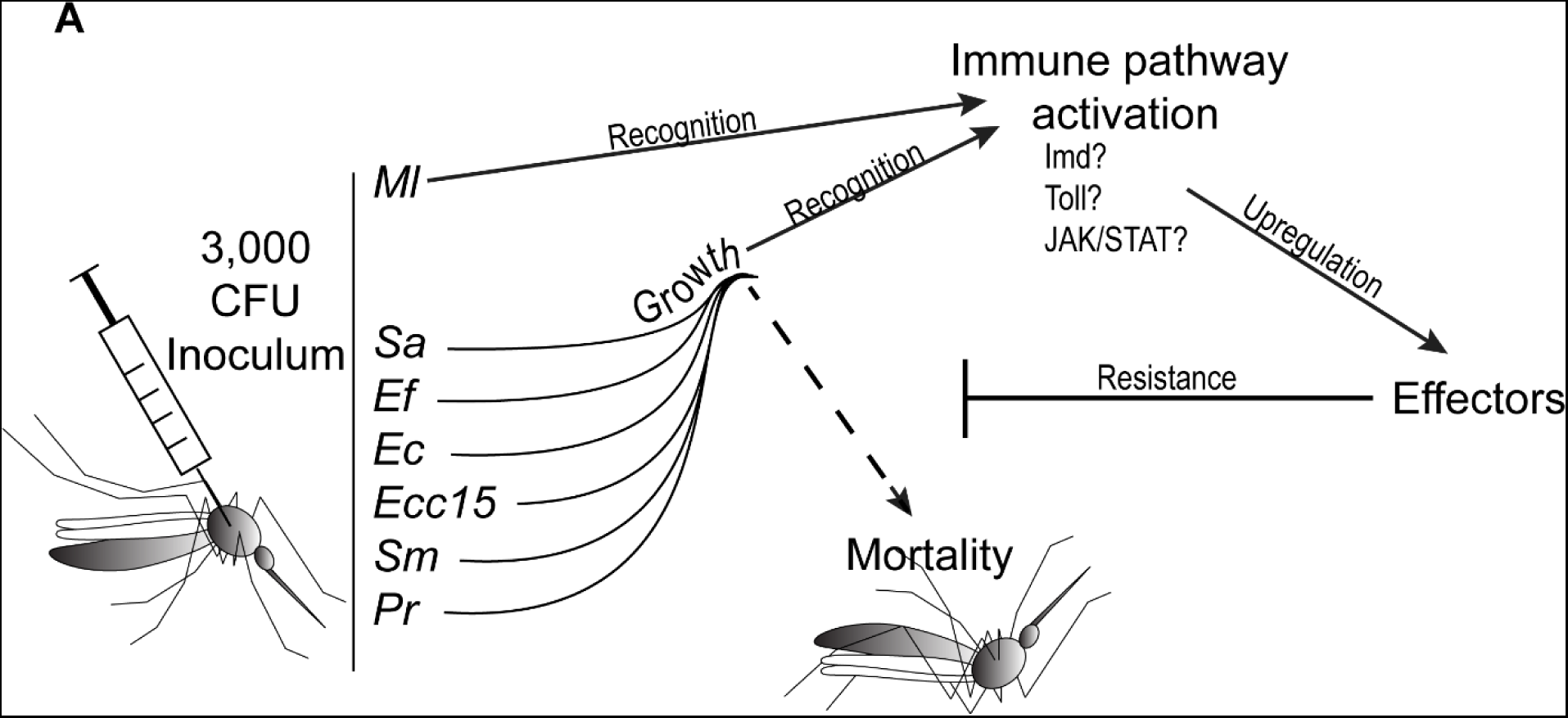
A scheme of the proposed interactions between bacterial pathogens, immune pathways, effector expression and mortality in challenged mosquitoes.

**Fig 6 S1:**
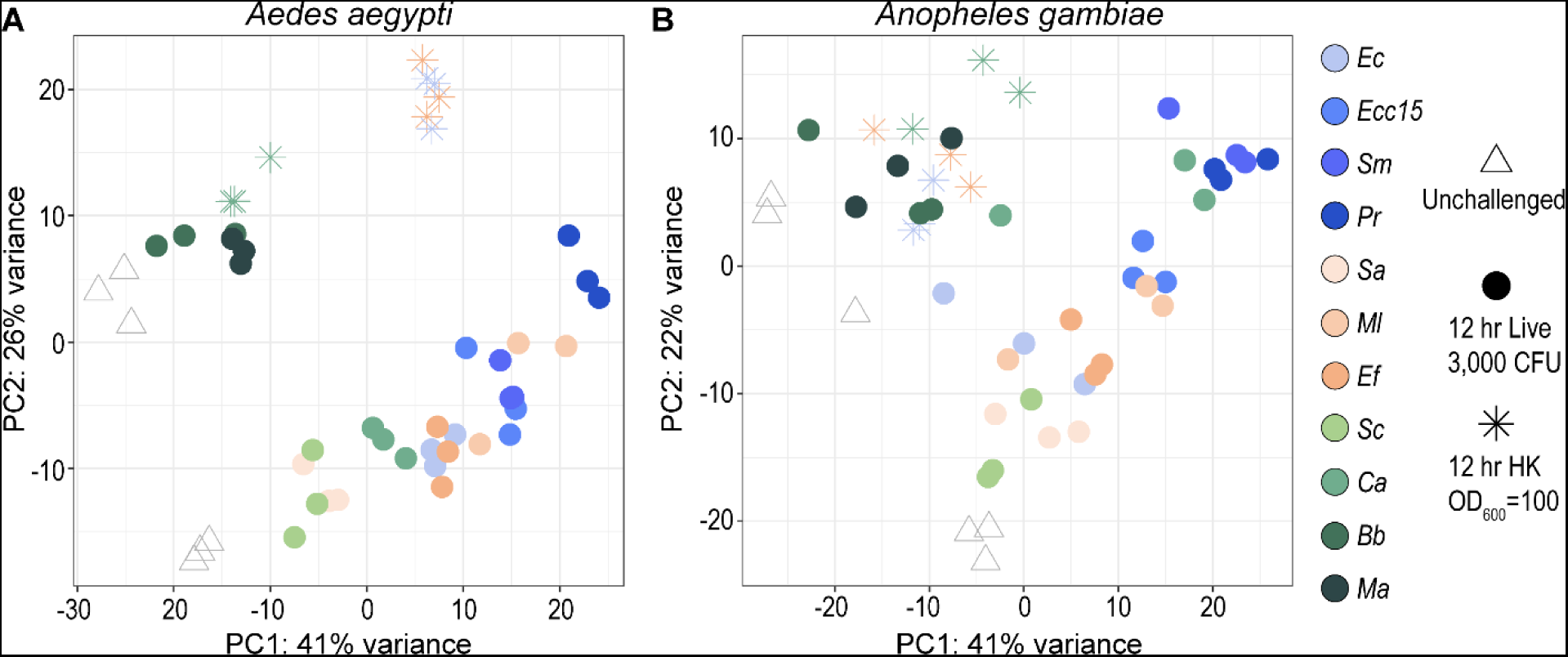
Principal component analyses combining transcriptomic data from two experiments in *Aedes aegypti* and *Anopheles gambiae*. Principal component analyses (PC1 vs PC2) of transcriptomes from *Ae. aegypti* (**A**) and *Anopheles gambiae* (*s.l.*) (**B**) combining data from two separate RNAseq experiments. From experiment #1: unchallenged (UC) mosquitoes and mosquitoes challenged with *Escherichia coli* (*Ec*), *Erwinia carotovora carotovora 15* (*Ecc15*), *Serratia marcescens* type strain (*Sm*), *Providencia rettgeri* (*Pr*), *Staphylococcus aureus* (*Sa*), *Micrococcus luteus* (*Ml*), *Enterococcus faecalis* (*Ef*), *Saccharomyces cerevisiae* (*Sc*), and *Candida albicans* (*Ca*). From experiment #2: unchallenged (UC) mosquitoes, and mosquitoes challenged with *Beauveria bassiana* (*Bb*), *Metarhizium anisopliae* (*Ma*), and concentrated (OD_600_ = 100) heat-killed (HK) *Ec*, *Ef*, and *Ca*; all data are from the 12-hour timepoint.

**Fig 6 S2:**
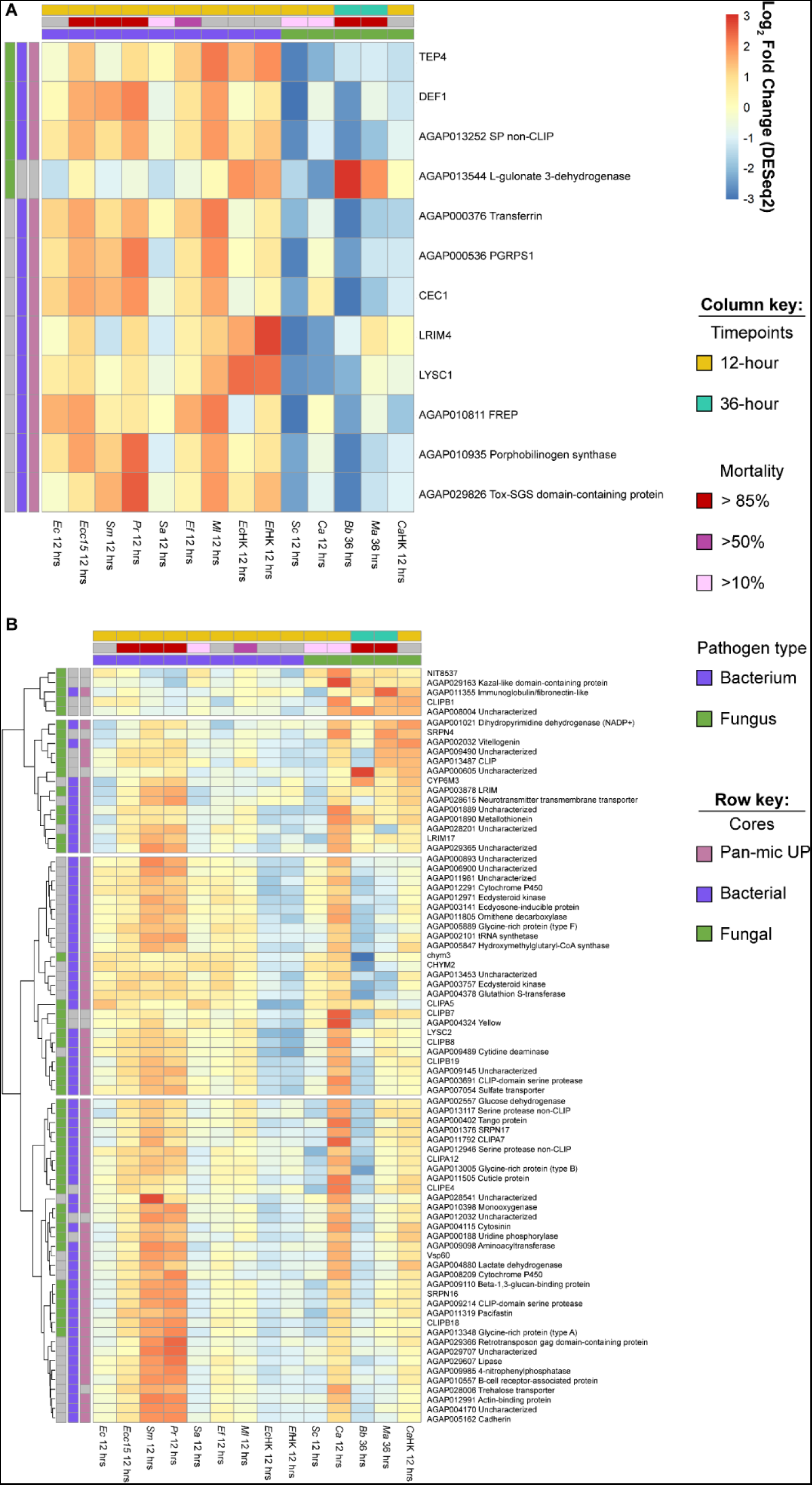
Heatmaps of expression in *Anopheles gambiae*. Heatmaps of log_2_fold-change in expression following challenge with *Escherichia coli* (*Ec*), *Erwinia carotovora carotovora 15* (*Ecc15*), *Serratia marcescens* type strain (*Sm*), *Providencia rettgeri* (*Pr*), *Staphylococcus aureus* (*Sa*), *Micrococcus luteus* (*Ml*), *Enterococcus faecalis* (*Ef*), *Saccharomyces cerevisiae* (*Sc*), *Candida albicans* (*Ca*), *Beauveria bassiana* (*Bb*), *Metarhizium anisopliae* (*Ma*), or with heat-killed *Ec* (*Ec*HK), heat-killed *Ef* (*Ef*HK), heat-killed *Ca* (*Ca*HK) at a concentration of OD_600_ = 100. (**A**) Genes belonging to the *An. gambiae* (*s.l.*) fungal and/or bacterial cores that are also differentially higher expressed in mosquitoes challenged with heat-killed bacteria (*Ec*HK and *Ef*HK replicates combined) versus *Ca*HK at 12 hours post-challenge. (**B**) Genes belonging to the *An. gambiae* fungal and/or bacterial cores that are also differentially higher expressed in mosquitoes challenged with *Ca*HK versus heat-killed bacteria. Criteria for differential expression are ≥ 1.5x fold-change and *padj* < 0.05, as calculated by DESeq2.

**Fig 6 S3:**
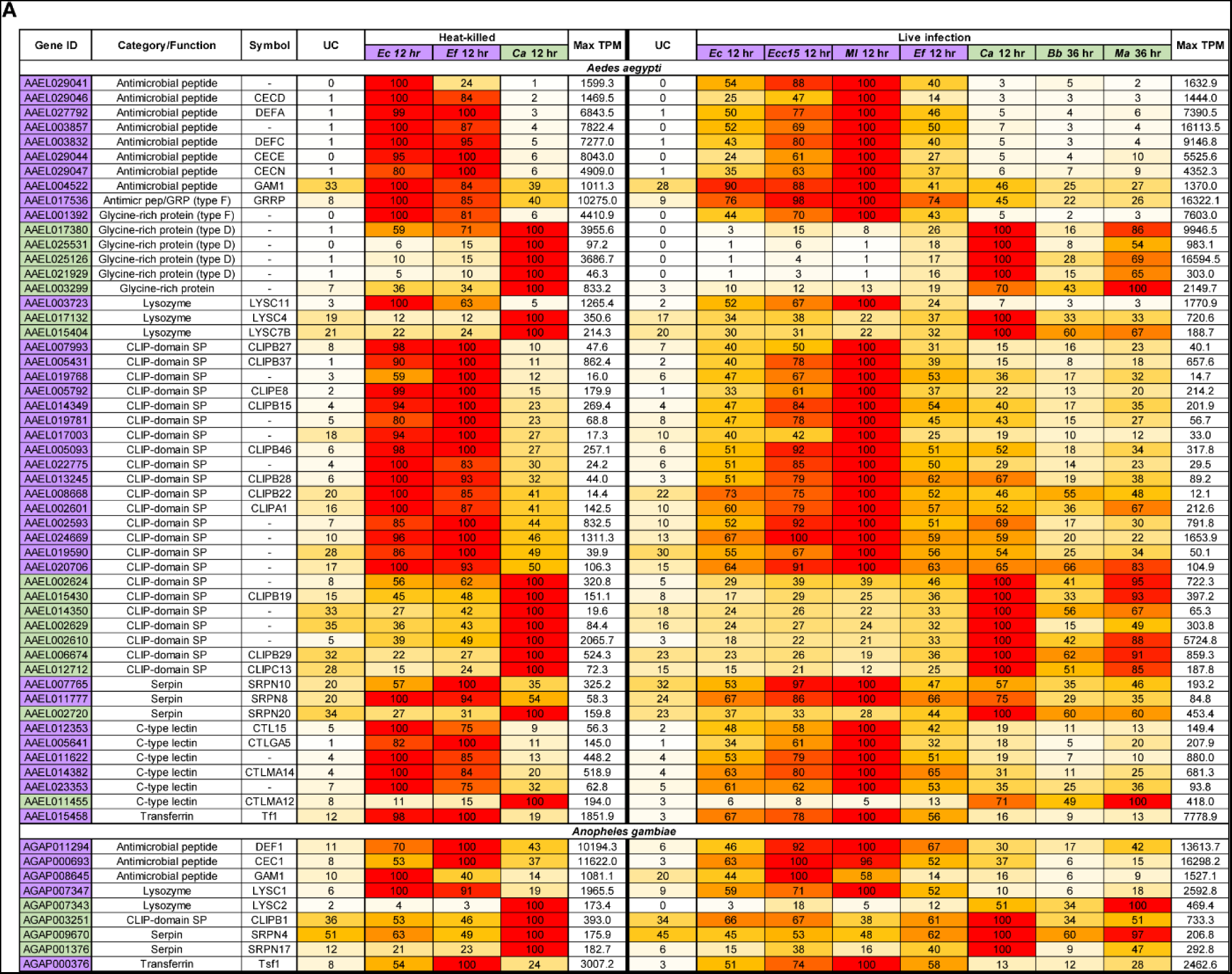
Selected genes in *Aedes aegypti* showing bacteria-specific or fungus-specific patterns of expression following heat-killed challenge or live infection. The left-hand side compares the expression of genes in unchallenged mosquitoes (UC) and mosquitoes challenged with heat-killed *Ec* (*Ec*HK), heat-killed *Ef* (*Ef*HK), or heat-killed *Ca* (*Ca*HK); all data from the second RNAseq experiment. The right-hand side compares unchallenged (UC), and challenged with live *Escherichia coli* (*Ec*), *Erwinia carotovora carotovora 15* (*Ecc15*), *Serratia marcescens* type strain (*Sm*), *Providencia rettgeri* (*Pr*), *Micrococcus luteus* (*Ml*), *Enterococcus faecalis* (*Ef*), *Candida albicans* (*Ca*) (from the first RNAseq experiment) and *Beauveria bassiana* (*Bb*) and *Metarhizium anisopliae* (*Ma*) (from the second RNAseq experiment). Within each comparison data are scaled to show relative expression so that the condition with the highest expression (shown in the max TPM column) is scored at ‘100’ and all lower expression values are expressed as a percentage of 100. Purple-labeled genes show greater responsiveness to bacterial challenges, while green-labeled genes show greater responsiveness to fungal challenges.

**Fig 6 S4:**
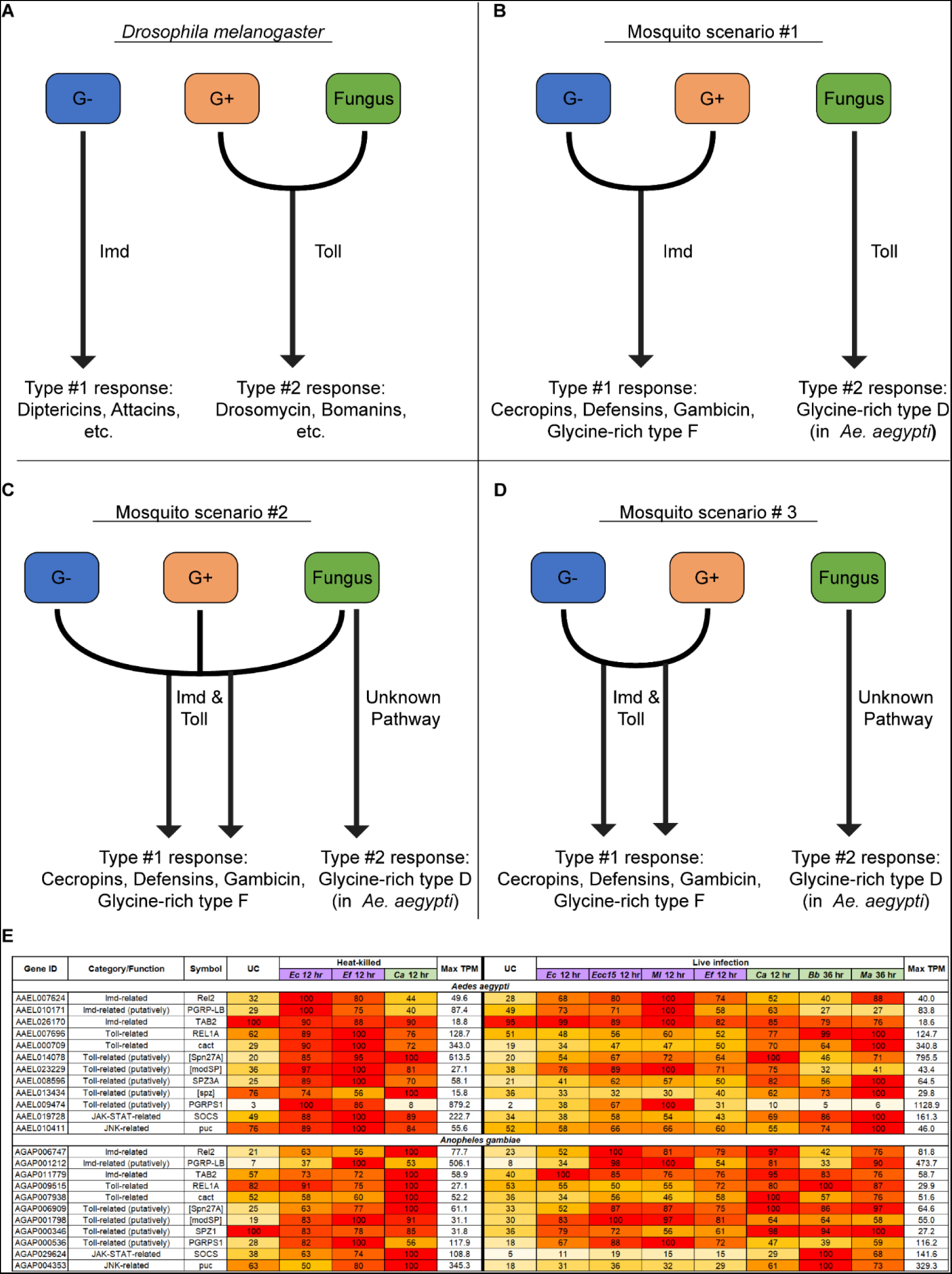
Models for the control of specific transcriptional responses to bacterial and fungal infection. (**A**) A model depicting the canonical specializations of the Imd and Toll pathways in *Drosophila melanogaster*. (**B**-**D**) Models depicting scenarios for the control of the transcriptional response to Gram-negative, Gram-positive, and fungal pathogens in mosquitoes. (**E**) A table displaying the expression patterns of genes associated (or putatively associated) with Imd, Toll, JAK-STAT, and JNK. The left-hand side compares the expression of genes in unchallenged mosquitoes (UC) and mosquitoes challenged with heat-killed *Ec* (*Ec*HK), heat-killed *Ef* (*Ef*HK), or heat-killed *Ca* (*Ca*HK); all data from the second RNAseq experiment. The right-hand side compares unchallenged (UC), and challenged with live *Escherichia coli* (*Ec*), *Erwinia carotovora carotovora 15* (*Ecc15*), *Serratia marcescens* type strain (*Sm*), *Providencia rettgeri* (*Pr*), *Micrococcus luteus* (*Ml*), *Enterococcus faecalis* (*Ef*), *Candida albicans* (*Ca*) (from the first RNAseq experiment) and *Beauveria bassiana* (*Bb*) and *Metarhizium anisopliae* (*Ma*) (from the second RNAseq experiment). Within each comparison data are scaled to show relative expression so that the condition with the highest expression (shown in the max TPM column) is scored at ‘100’ and all lower expression values are expressed as a percentage of 100.

